# Multi-environment genome-wide association mapping of culm morphology traits in barley

**DOI:** 10.1101/2022.03.30.486427

**Authors:** G. Bretani, S. Shaaf, A. Tondelli, L. Cattivelli, S. Delbono, R. Waugh, W. Thomas, J. Russell, H. Bull, E. Igartua, A. Casas, P. Gracia, R. Rossi, A. Schulman, L. Rossini

## Abstract

In cereals with hollow internodes, lodging resistance is influenced by morphological characteristics such as internode diameter and culm wall thickness. Despite their relevance, knowledge of the genetic control of these traits and their relationship with lodging is lacking in temperate cereals such as barley. To fill this gap, we developed an image-analysis based protocol to accurately phenotype culm diameter and culm wall thickness across 261 barley accessions. Analysis of culm trait data collected from field trials in 7 different environments revealed genetic control as supported by high heritability values, as well as genotype-by-environment interactions. The collection was structured mainly according to row-type, which had a confounding effect on culm traits as evidenced by phenotypic correlations. In addition, culm traits showed strong negative correlations with lodging but weak correlations with plant height across row-types, indicating the possibility of improving lodging resistance independent of plant height. Using 50k iSelect SNP genotyping data, we conducted multi-environment genome-wide association studies using mixed model approach across the whole panel and row-type subsets: we identified a total of 192 QTLs for the studied traits, including subpopulation-specific QTLs and several main effect loci for culm traits showing negative effects on lodging without impacting plant height. Providing first insights into the genetic architecture of culm morphology in barley and the possible role of candidate genes involved in hormone and cell wall related pathways, this work supports the potential of loci underpinning culm features to improve lodging resistance and increase barley yield stability under changing environments.

**One-sentence summary:** Genetic analysis of a diverse collection of European barleys reveals genomic regions underpinning stem morphological features associated with lodging resistance.

## Introduction

Selection of desired plant architecture traits has represented a driving force in crop domestication and breeding. In cereals, one of the most paradigmatic examples is offered by the widespread introduction of semi-dwarfing genes in the modern varieties of the Green Revolution. When high fertilizer inputs were applied, traditional varieties elongated and lodged, i.e. fell over leading to major losses in grain yields (Islam et al., 2007; Berry, 2013; Piñera-Chavez et al., 2016). To avoid this problem, breeders developed new semi-dwarf varieties with reduced plant height and sturdy stems, improving lodging resistance and crop production (Khush, 2001; Chandler and Harding, 2013). Several semi-dwarfing genes are involved in the pathways of gibberellins (GA) and brassinosteroids (BR), phytohormones which play a major role in stem elongation (Sasaki et al., 2002; Kuczyńska et al., 2013). Examples of alleles deployed in breeding include loss-of-function mutations of the rice (*Oryza Sativa*) *semidwarf* (*SD1*) locus encoding a *OsGA20ox2* involved in GA biosynthesis (Sasaki et al., 2002). In wheat (*Triticum aestivum* L), mutants of *Reduced Height-1* (*Rht*) genes are responsible for the expression of mutated forms of DELLA GA signalling repressor proteins (Peng et al., 1999). In barley (*Hordeum vulgare*), *semi-dwarf 1* (*sdw1*) and *semi-brachytic 1* (*uzu1*) mutant alleles were widely used in breeding programs (Kuczyńska et al., 2013; Xu et al., 2017). Barley *Sdw1* encodes a GA 20-oxidase (like rice SD1), while a missense mutation in the BR receptor gene *HvBRI1* causes the *uzu* phenotype (Chono et al., 2003; Kuczynska and Wyka, 2011). Despite providing yield gains, some semi-dwarfing alleles have been associated to negative pleiotropic effects such as temperature sensitivity, late flowering and reduced grain quality (Rajkumara, 2008; Okuno et al., 2014).

Changes in climatic conditions are predicted to increase intensity and frequency of storms, hail and heavy rains (Lobell et al., 2011), the major causes of lodging impacting crop productivity (Berry and Spink, 2012; Berry, 2013). In cereals such as rice, wheat and barley, the stem or culm consists of alternating solid nodes and hollow internodes. Three different types of lodging are known: culm bending, culm breaking and root lodging (Hirano et al., 2017a). Breaking-type lodging is more serious than bending type because bent culms are still able to transport photosynthetic assimilates from the leaves to the panicles, which is necessary for plant recovery and grain filling. Since cereal height cannot be reduced below a certain point, improvement of lodging resistance and therefore yield requires identification and use of other important traits (Dawson et al., 2015; Hirano et al., 2017a; Shah et al., 2019).

Barley is one of the most important crops worldwide. Due to its intrinsic plasticity and adaptability, barley can be cultivated in areas not suited to maize and wheat, especially where the climatic conditions are cool and/or dry. Barley varieties can be divided into two-row and six-row types. In two-row barley, the central spikelet of each triplet on the rachis is fertile, while the other two are reduced and do not develop. Mutations of the *VRS1* gene determine the fertility of these lateral spikelets to produce six-row barleys (Komatsuda et al., 2007), and have pleiotropic effects on a number of morphological traits (Liller et al., 2015).

Barley production can be lowered from 4 to 65% by lodging (Jedel and Helm, 1991; Sameri et al., 2009). While agricultural practices play an important role (Cai et al., 2019), the occurrence of culm bending/breaking lodging events is determined mainly by two factors: 1) the force exerted on the culm (e.g. wind-induced forces or panicle weight) (Pinthus, 1974) and 2) the mechanical resistance of the stem determined by composition and morphology (Samadi et al., 2019).

For example, in cereals with hollow internodes such as barley and rice, lodging resistance is influenced by morphological characteristics such as internode diameter and culm wall thickness (Samadi et al., 2019; Zhang et al., 2020). Wider culm diameter and thickness were shown to improve lodging resistance e.g. in wheat (Zuber et al., 1999). Also a stronger culm may help to improve yield by allowing increased nutritional inputs. Despite the relevance of these traits, knowledge of the genetic control of culm diameter and culm wall thickness is limited to few studies in rice. A rice mutant with larger stem diameter and thickness called *smos1* (*small organ size*) exhibits altered cell wall composition and is less prone to lodging (Hirano et al., 2014). The *SMOS1* gene encodes an APETALA2 (AP2)-type transcription factor (Aya et al., 2014; Hirano et al., 2014) that interacts with a GRAS transcription factor encoded by *SMOS2/DLT* to mediate cross-talk between auxin and BR signalling and regulate various culm morphology features (Hirano et al., 2017b). In rice cultivar Habataki, a variety with improved yield and large culms, two QTLs associated with culm architecture: *STRONG CULM1* (*SCM1*) and *SCM2*/*APO1* (*ABERRANT PANICLE ORGANIZATION1*) were respectively identified on chromosome 1 and chromosome 6 (Ookawa et al., 2010). Two additional *SCM* loci were identified from the high yielding and lodging resistant cultivar Chugoku 117, including *SCM3* which was shown to be allelic to the rice *TEOSINTE BRANCHED1* (*OsTB1*)/*FINE CULM1* (*FC1*) gene (Minakuchi et al., 2010; Yano et al., 2015; Cui et al., 2020). Recently, the mediator subunit gene *OsMED14_1* was uncovered as a new player in culm and lateral organ development through *NARROW LEAF1* (*NAL1*) gene regulation (Malik et al., 2020).

The lack of efficient and accurate phenotyping protocols has been a limiting factor in further genetic dissection of culm architecture for example through exploration of wider genetic diversity in germplasm collections. In this context, different solutions emerged in recent years relying on high-throughput phenotyping methods based on the use of new image analysis tools with advanced software and special platforms (Agnew et al., 2017).

So far little is known about the genetic architecture underlying barley culm development and morphology. Aims of this work were to explore natural genetic diversity for culm architecture traits in barley, analyze their correlations with plant height, lodging and phenology, and identify associated genomic regions and candidate genes through multi-environment genome-wide association studies (GWAS) on a collection of 261 European accessions. To these ends, we developed an image-analyses based protocol to accurately phenotype culm diameter and culm wall thickness and integrated the resulting data with genome-wide marker data from 50k SNP iSelect genotyping (Bayer et al., 2017).

## Results

### Diversity, population structure and linkage disequilibrium of the barley panel

The barley panel considered in the present study is a collection representing the diversity of European barley from the 20th century and was chosen based on previous geographic and genetic diversity analysis (Tondelli et al., 2013). This panel was supplemented with 57 six-row and five two-row Spanish landraces representing the ecogeographic diversity of barley cultivation in the Iberian Peninsula. Eight of the 269 genotypes did not match with their phenotypes and were discarded from the analyses resulting in a total of 261 barley cultivars and landraces comprising 165 two-row and 96 six-row barleys being considered in this study (Supplemental Table S1). The 50k SNP iSelect genotyping of the collection yielded a set of 33342, 26262, and 27583 polymorphic markers for the whole, two-row, and six-row panel, respectively (Supplemental Table S2; Supplemental Figure S1).

Genetic structure of the panel was investigated using Principal Component Analysis (PCA) on a pruned subset of markers to reduce the effect of linkage disequilibrium (LD) on population structure. PCA indicated the first two PC scores explained, respectively, 13% and 8.5% of total variation (Supplemental Figure S2 A). The first PC could distinguish six-row from two-row barleys, while the second PC axis was attributed to separation of landraces from cultivars within six-row barleys. In addition, PCA revealed the wider level of genetic variation within six-row barleys, although the proportion of two-row barleys was higher in the panel.

As a prerequisite for GWAS, LD was calculated for each chromosome using the squared correlation coefficient between marker pairs, r^2^, after correcting for genomic relatedness. The LD decay was visualized by plotting r^2^ values against the physical distance in Mb. Considerable variation was observed across the genome among whole panel and row-type subsets, reflecting breeding history and effect of selection (Supplemental Figure S2 B-D). The level of LD decay in the two-row panel at the critical r^2^ threshold was higher (LD = 1.4 Mb) compared to LD decay observed within the six-row panel (LD = 0.6 Mb), with slightly higher LD in the whole panel (LD = 0.8 Mb).

### Phenotypic variation, trait heritability and correlations

The barley collection was grown under field conditions in seven environments including four locations and two years, 2016 and 2017 (Supplemental Table S3). Field sites were chosen to represent contrasting environments in southern Europe (Italy, CREA; Spain, CSIC) and northern Europe (Scotland, JHI; Finland, LUKE). Regarding culm traits, we focused on culm features reported in the literature as critical for lodging resistance in hollow cereals (Ookawa et al., 2010). Because of the great plasticity of the first internode, we decided to focus on the second basal internode as a critical point for lodging resistance and a good descriptor of culm characteristics (Pinthus, 1974; Berry et al., 2004). For all trials outer culm diameter (OD), inner culm diameter (ID), culm thickness (TH) were quantified using a newly developed image analysis-based protocol (Figure 1; Supplemental Methods S1). In order to investigate the correlations between culm traits and some agronomic traits, we also included heading (HD), plant height (PH), and lodging (LG) (Supplemental Table S4). We further derived section modulus (SM), the ratio between OD and TH (herewith designated as stiffness, ST) and the ratio between OD and PH (stem index, SI) as indexes reflecting physical strength of the culm (Supplemental Table S4; Mulsanti et al., 2018; Sowadan et al., 2018). For trial CSIC16 it was not possible to collect lodging data. The best linear unbiased predictions (BLUEs) were calculated for the downstream analyses.

**Figure 1.**
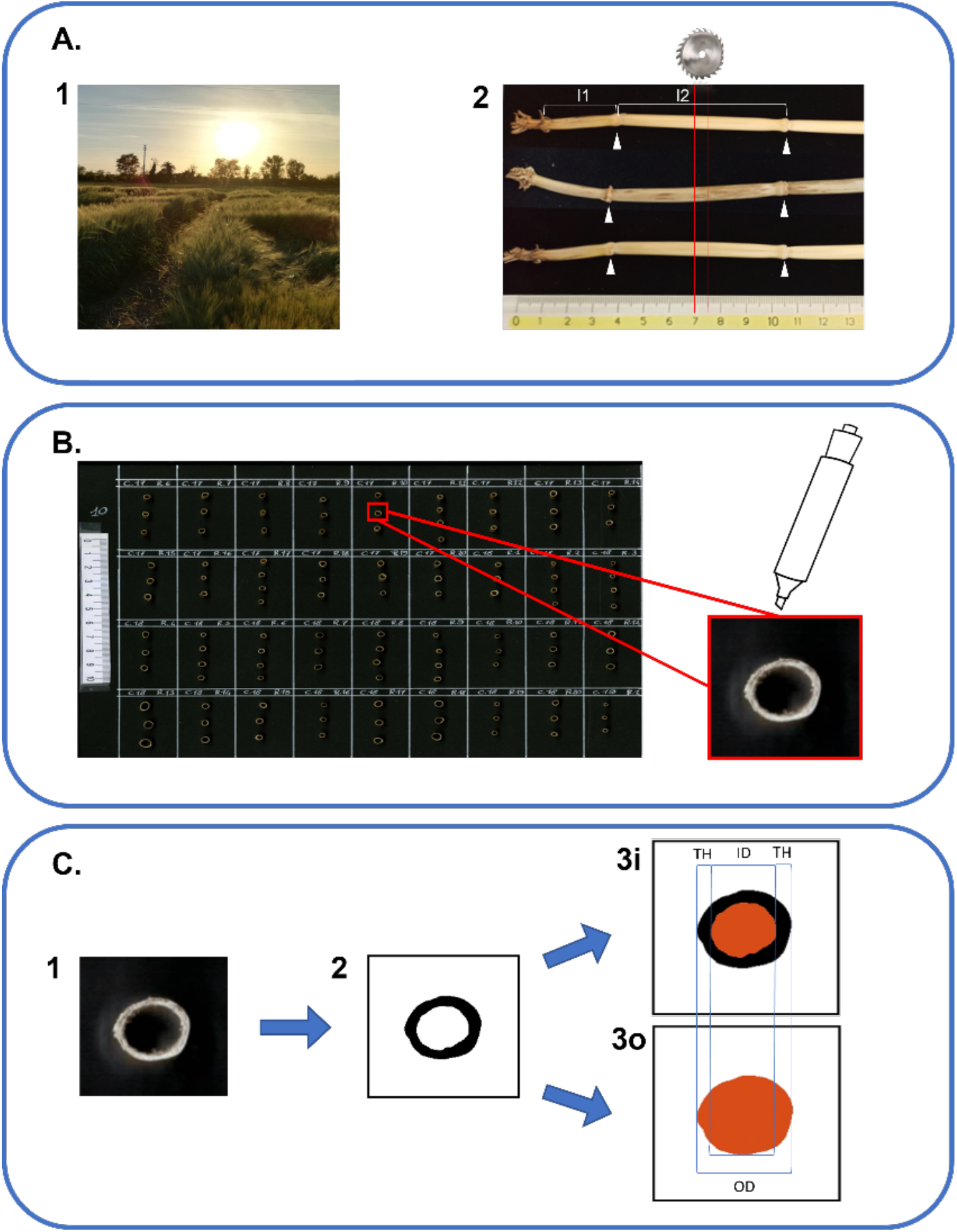
Workflow of phenotyping protocol for culm morphology traits. **A.1**) Barley specimens were gathered when plants reached Zadoks stage 90 (grain ripening). Three random plants were collected from each plot. **A.2**) Samples were cleaned and the main culm was selected for each plant. The first internode (I1) was identified as the most basal internode ≥ 1cm. The second internode (I2) was the one immediately above (white arrowheads indicate the positions of flanking nodes). Five mm tall sections from the center of I2 (red lines) were obtained using a dedicated circular saw. **B**) Sections were attached to black A4 cardboaord with superglue and organized on the cardboard following the plot order in the field. The upper part of each section was highlighted with a bright white marker in order to enhance the contrast with the blackboard. **C.1**) Cardboards with I2 sections were scanned using a flat office scanner to obtain 300 dpi color images. **C.2**) Using the software ImageJ with a dedicated macro the I2 section images were converted to black and white images. **C.3i**) ImageJ software was able to isolate and measure the medullary cavity of the culm (in red). **C.3o**) ImageJ software was used to isolate and measure the external outline (in red). ID, inner diameter, OD, outer diameter, TH, thickness were derived from images 3i and 3o according to formulas in Supplemental Table S4.

The single and across environment means, standard deviations (SDs), ranges, minimum, and maximum values are indicated in Supplemental Table S5. Considerable phenotypic variation was present both within and across environments. In general, for all traits higher mean values were observed for Southern environments. CSIC16 had the highest values for almost all culm traits in the whole panel, and both two-row and six-row panels. Highest values for HD were recorded in the CREA17 trial, while CREA16 had the highest mean value for PH in the whole panel and also two-row and six-row panels.

Heritability values were calculated both in single and combined environments in the whole panel and both two-row and six-row subsets (Table 1; Supplemental Methods S3). In most environments, analysis of variance correcting for field trends i.e. the correlation between residuals from neighboring plots using the first-order autoregressive model (AR1), improved the precision compared to base model fitting. High heritability values (>50%) were obtained for most traits except for TH and ST, although these traits showed improved heritability in the combined environment analysis compared to single environment. Heritability estimates varied among environments indicating the presence of heterogeneity of genotype variance due to genotype × environment interactions. This was especially evident for TH and ST due to their relatively low heritability values.

**Table 1.** Estimates of broad-sense heritability for culm morphological traits in single and across environments. Heritabilities with less than moderate valule are indicated in bold.

| Env | Factor <sup>a</sup> | Residual <sup>b</sup> | Panel | HD | PH | OD | ID | TH | SM | ST | SI |
| --- | --- | --- | --- | --- | --- | --- | --- | --- | --- | --- | --- |
| CREA16 | Gen+Rep+Col+Row | AR1(Row) : AR1(Col) | Whole panel | 98.75 | 90.00 | 80.48 | 81.12 | 53.31 | 75.97 | 59.79 | 87.81 |
|  |  |  | 2-row | 96.79 | 82.64 | 65.44 | 69.33 | **27.95** | 53.13 | 53.36 | 83.82 |
|  |  |  | 6-row | 99.32 | 87.43 | 86.14 | 87.90 | 55.09 | 83.87 | 67.24 | 90.21 |
| CREA17 | Gen+Rep+Col | ID(Row) : AR1 (Col) | Whole panel | 94.02 | 85.76 | 73.84 | 65.35 | 67.01 | 75.59 | 51.75 | 86.42 |
|  |  |  | 2-row | 87.47 | 82.84 | 70.22 | 64.19 | **47.37** | 67.04 | 56.27 | 82.89 |
|  |  |  | 6-row | 95.89 | 88.77 | 79.01 | 70.85 | 77.18 | 82.85 | **39.37** | 90.70 |
| CSIC16 | Gen+Rep+Col+Row | ID(Row) : ID (Col) | Whole panel | 97.13 | 94.84 | 79.58 | 71.65 | 77.51 | 81.93 | 52.39 | - |
|  |  |  | 2-row | 94.39 | 90.15 | 75.29 | 67.85 | 54.87 | 73.45 | **32.22** | - |
|  |  |  | 6-row | 98.35 | 92.59 | 81.16 | 76.19 | 78.27 | 85.01 | 64.09 | - |
| CSIC17 | Gen+Rep+Col+Row | AR1(Row) : AR1(Col) | Whole panel | 94.17 | 93.11 | 87.39 | 81.67 | 78.77 | 86.67 | 72.08 | 92.81 |
|  |  |  | 2-row | 86.88 | 85.18 | 79.64 | 78.16 | **29.82** | 69.25 | 70.15 | 89.71 |
|  |  |  | 6-row | 96.51 | 92.85 | 85.95 | 82.89 | 78.08 | 87.43 | **48.81** | 94.76 |
| JHI16 | Gen+Rep+Col+Row | AR1(Row) : AR1(Col) | Whole panel | 95.62 | 93.85 | 89.54 | 87.51 | 81.98 | 90.00 | 73.07 | 92.15 |
|  |  |  | 2-row | 90.85 | 93.48 | 77.53 | 68.67 | 70.85 | 72.26 | 55.07 | 91.87 |
|  |  |  | 6-row | 87.97 | 90.45 | 88.57 | 89.22 | 76.56 | 89.35 | 74.01 | 91.14 |
| JHI17 | Gen+Rep+Col+Row | AR1(Row) : AR1(Col) | Whole panel | 93.37 | 93.13 | 84.64 | 78.33 | 53.32 | 86.75 | **47.19** | 85.73 |
|  |  |  | 2-row | 83.68 | 92.78 | 69.05 | 63.14 | **20.85** | 59.37 | **35.96** | 85.22 |
|  |  |  | 6-row | 86.67 | 93.60 | 85.45 | 78.39 | 67.04 | 88.21 | **41.48** | 81.15 |
| LUKE17 | Gen+Rep+Col+Row | ID(Row) : ID (Col) | Whole panel | 95.39 | 96.74 | 84.74 | 84.87 | 61.62 | 84.75 | 60.60 | 92.04 |
|  |  |  | 2-row | 91.05 | 93.33 | 57.80 | 57.34 | **41.01** | 50.03 | **43.83** | 91.04 |
|  |  |  | 6-row | 96.44 | 95.74 | 84.62 | 86.14 | 54.34 | 84.50 | 62.32 | 89.19 |
| **Combined Environments** |  |  | Whole panel | 93.10 | 91.42 | 89.21 | 86.98 | 81.52 | 84.77 | 70.49 | 91.91 |
|  |  |  | 2-row | 90.43 | 94.26 | 83.31 | 81.37 | 57.81 | 72.38 | 57.98 | 92.48 |
|  |  |  | 6-row | 89.95 | 81.90 | 90.18 | 88.98 | 68.61 | 85.49 | 75.11 | 91.58 |

**a: (Gen):** random genotype effect; **(Rep):** Random replicate effect; **(Row):** Random row effect; **(Col):** random column effect; **ID (Row):ID(Col):** two dimensional independent error structure; **ID(Row):AR1(Col):** One dimensional correlated error for columns with first-order autoregressive process ; **AR1(Row):ID(Col):** one dimensional correlated error for rows with first-order autoregressive process ; **AR1(Row):AR1(Col):** two dimensional correlated error with first-order auto regressive process.

We further compared phenotypic means according to row-type and germplasm source as these were important factors shaping population structure within the panel (Supplemental Figures S2 A, S3). Results showed that two-row landraces and six-row cultivars had latest and earliest heading, respectively, in southern trials, while two-row cultivars were latest heading in northern trials. In these comparisons however, it should be noted that only 6 two-row landraces were included in our collection, all from Spain, providing limited representation of this category. PH was highly variable across environments and was mainly highest for six-row landraces in southern trials, but this was highest for mainly two-row landraces in northern locations. LG was lowest in all environments in two-row cultivars and highest in six-rowed landraces. For culm morphology, six-row cultivars showed highest values of OD, ID, SM, SI, and TH, whereas two-row landraces were lowest almost in all environments. ST was however highly variable both within and between northern and southern trials. Based on phenotypic values obtained from combined analysis of environments, higher values were observed for culm morphological traits in the cultivar gene pool, especially in six-row cultivars, but two-row cultivars were on average less susceptible to lodging. Generally, landraces showed higher values for PH and HD.

Together, these analyses show that our germplasm panel harbors significant genetic variation for culm-related traits and suggest the existence of complex genotype × environment interactions. The obtained datasets provide and ideal starting point for investigating the genetic architecture of barley culm morphology under contrasting environmental conditions.

In order to gain insight into the relationships among different traits, pairwise correlations were calculated based on phenotype values estimated both within single and combined analysis of environments (Fig.2 A, Supplemental Figures S4-S6). Germplasm source and row-type were also considered to study their relationship with the different traits. These values were also calculated within two-row and six-row panels to control for row-type. In the whole panel, row-type showed positive correlations with LG, PH and culm morphological traits, but negatively correlated with ST, SI, and HD. Germplasm source (cultivars coded as presence) had negative correlations with PH, TH, and LG and positive correlations with OD, ID, SI, and ST, meaning that cultivars were shorter and less prone to lodging with larger culm diameter compared to landraces. However, correlation between germplasm source and HD was dependent on region with positive values in northern environments and negative values in southern sites. Results show that in the whole panel strong correlations were present between culm morphological traits. Similar results were also obtained in single environments (Supplemental Figures S4-S6). Except for TH, culm traits were negatively correlated with LG and HD, but positively correlated with PH. As expected, LG was positively correlated with PH. Taken together, correlation analyses on the whole panel show that in our collection six-row lines tended to have wider and thicker culms and were overall more prone to lodging compared to two-row. While a confounding effect of row-type may account for the relatively weak correlations between LG and culm diameter and thickness, it should be also noted that in our germplasm collection landraces are more represented in the six-row subset compared to the two-row subset: this may be a confounding factor contributing to observed differences between the row-type subsets.

**Figure 2.**
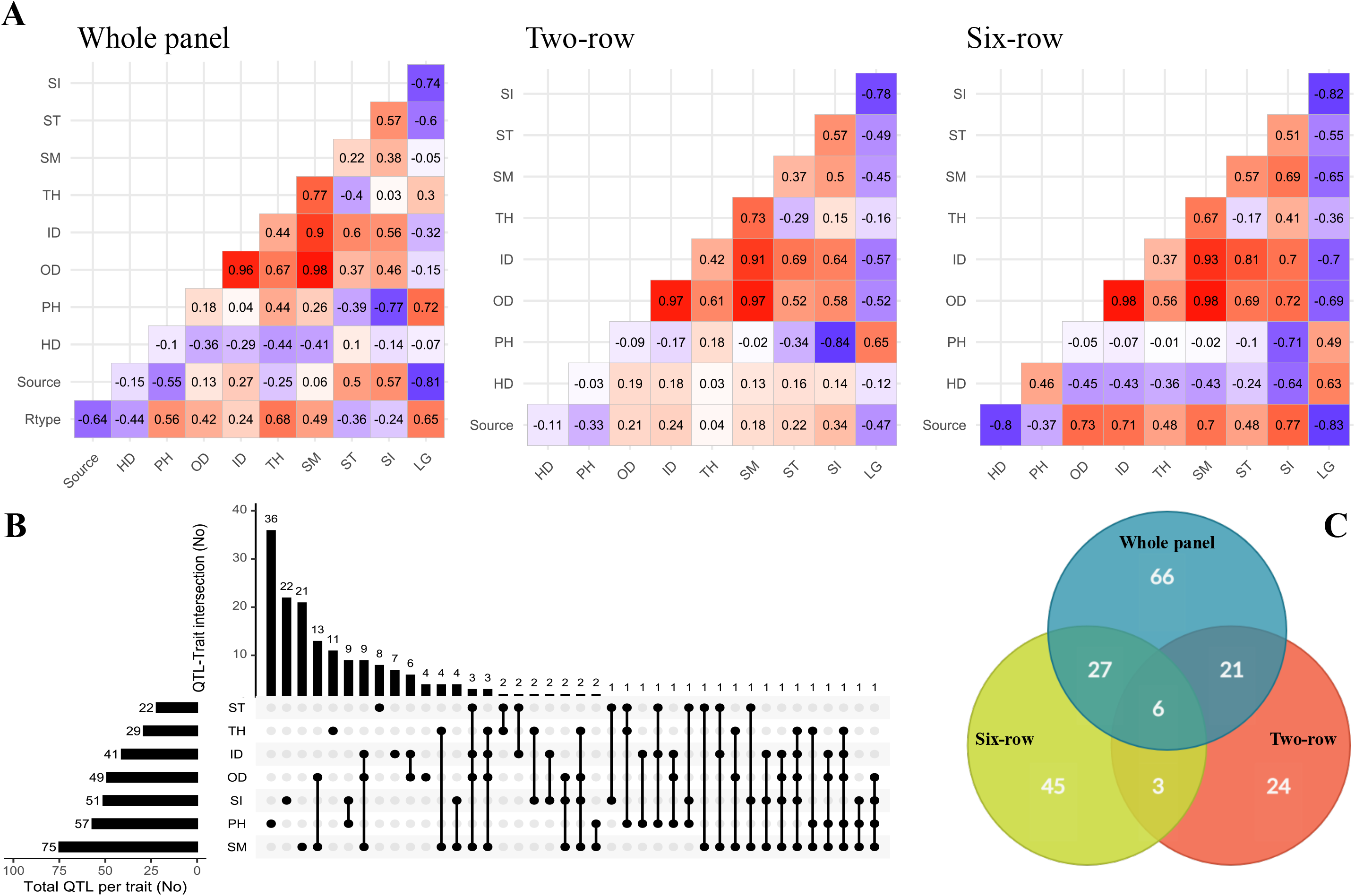
A) Pairwise phenotypic correlations between traits along with row type and germplasm sources within whole panel and row type groups based on means estimated across trials; B) UpSetR plot showing the overlap of the associated SNPs/loci for traits identified by GWAS; C) Venn diagram showing distribution of QTLs among whole panel and row type groups.

In order to explore the relationships between culm traits and lodging, excluding the effect of row-type, further analyses were conducted within row-type subsets.

In the two-row panel, correlations between culm traits were generally maintained and stronger negative correlations were observed between culm morphological traits and lodging. Some discrepancies were also observed compared to the whole panel, e.g. the negative relationship between TH and lodging in contrast to the positive correlation between these traits in the whole panel, which was possibly due to confounding effects from six-row landraces (thick culms and more prone to lodging). Furthermore, while positively correlated with lodging, PH was environment-dependent and did not show strong correlations with culm morphology, e.g. in southern environments the relationship was mainly weakly negative and in northern weakly positive (Supplemental Figure S5). HD was also mainly positively correlated with culm morphology.

In the six-row panel, culm morphological traits had the strongest interrelationships. HD was also in agreement with whole panel with stronger negative correlations with culm morphology, and in contrast to the two-row panel, it was positively correlated with lodging. PH had negatively weak relationship with culm traits with stronger positive correlations in northern trials and negative correlations in southern trials (Supplemental Figure S6).

Together, these results highlight the potential of culm morphological traits as interesting targets for improvement of lodging resistance in barley. In particular, the general lack of correlation within row-type subsets suggests that culm diameter is largely controlled by distinct genetic factors with respect to PH.

### Multi-environment genome-wide association mapping

We performed GWAS using multi-trait mixed model (MTMM) proposed for multi-trait or multi-environment association mapping to detect quantitative trait loci (QTLs) underlying culm morphological traits, incorporating kinship estimated from marker data and population structure using principal components (Korte et al., 2012). This method allows to identify five types of marker-trait associations: markers with main effects stable across environments (QM), markers with main but also significant interaction effects (QF), marker-by-environment interaction effects (QE), marker-by-location interaction effect (QL), and marker-by-year interaction effect (QY) (see Supplemental Methods S4 for more details). GWAS of multi-environment trials were performed for the whole panel and also for two-row and six-row subsets separately. The experiment-wise GWAS significance threshold was determined according to the actual number of independent SNP tests as estimated in Haploview software using the tagger function and the r^2^ threshold estimated from LD decay analysis. These threshold values were found to be -log_10_ (P) ≥ 4.94, -log_10_ (P) ≥ 4.75, and -log_10_ (P) ≥ 5.02 for the whole panel, two-row, and six-row panels, respectively. However, the p-values with -log_10_ (P) ≥ 4 were also retained as suggestive QTLs.

A total of 732 marker-trait associations were detected, and the associated SNPs with -log_10_ (P) ≥ 4 in close vicinity were grouped into a single QTL based on the average LD decay, due to variable LD blocks for individual chromosomes and thus a variable decay across the chromosomes (Supplemental Figure S2 B-D). This allowed us to converge marker-trait associations into 192 QTLs (93 single SNPs and 99 SNP clusters) across the whole, two-row and six-row panels (Supplemental Table S6). From these loci, 109 were trait-specific and the remaining were co-associated to at least two traits (Fig. 2B). PH with 36 QTLs and OD with four QTLs were the traits with the maximum and minimum number of specific QTLs. Most QTLs were co-associated between culm morphological traits. Among the highest number of co-associated QTLs, 13 QTLs were common between SM and OD, 9 QTLs between PH and SI, 9 QTLs between ID, OD and SM, and 6 QTLs were commonly associated with ID and OD. In agreement with a largely independent genetic control, the lowest number of co-associated QTLs were identified between PH and culm morphological traits. In addition, 66, 24, and 45 QTLs were specific to the whole panel, two-row, and six-row panels, respectively (Fig. 2C). Other QTLs were in common between at least two panels.

Co-association network analysis for the 192 QTLs revealed many co-association modules across the whole panel and the row-type sub-panels, each of which contained loci from one or more genomic regions distributed on different chromosomes (Fig. 3). The co-association module is a cluster of one or more loci that are connected by edges. The edges connecting two loci have similar associations with the phenotype with a distance below the threshold. Loci in different clusters are more dissimilar than to those in the same group and would not be connected by edges in a co-association module. In other words, associated nodes with edges appeared in close proximity, while weakly associated nodes appear far apart. One common feature that can be clearly derived from this visualization was that PH and SI were in closer proximity across all panels and nodes for culm morphological traits were closer together and far apart from PH. There were however some exceptions especially for ST and TH that exhibited higher dispersion. Another interesting observation is that loci with the same type of QTL effect appeared closer.

**Figure 3.**
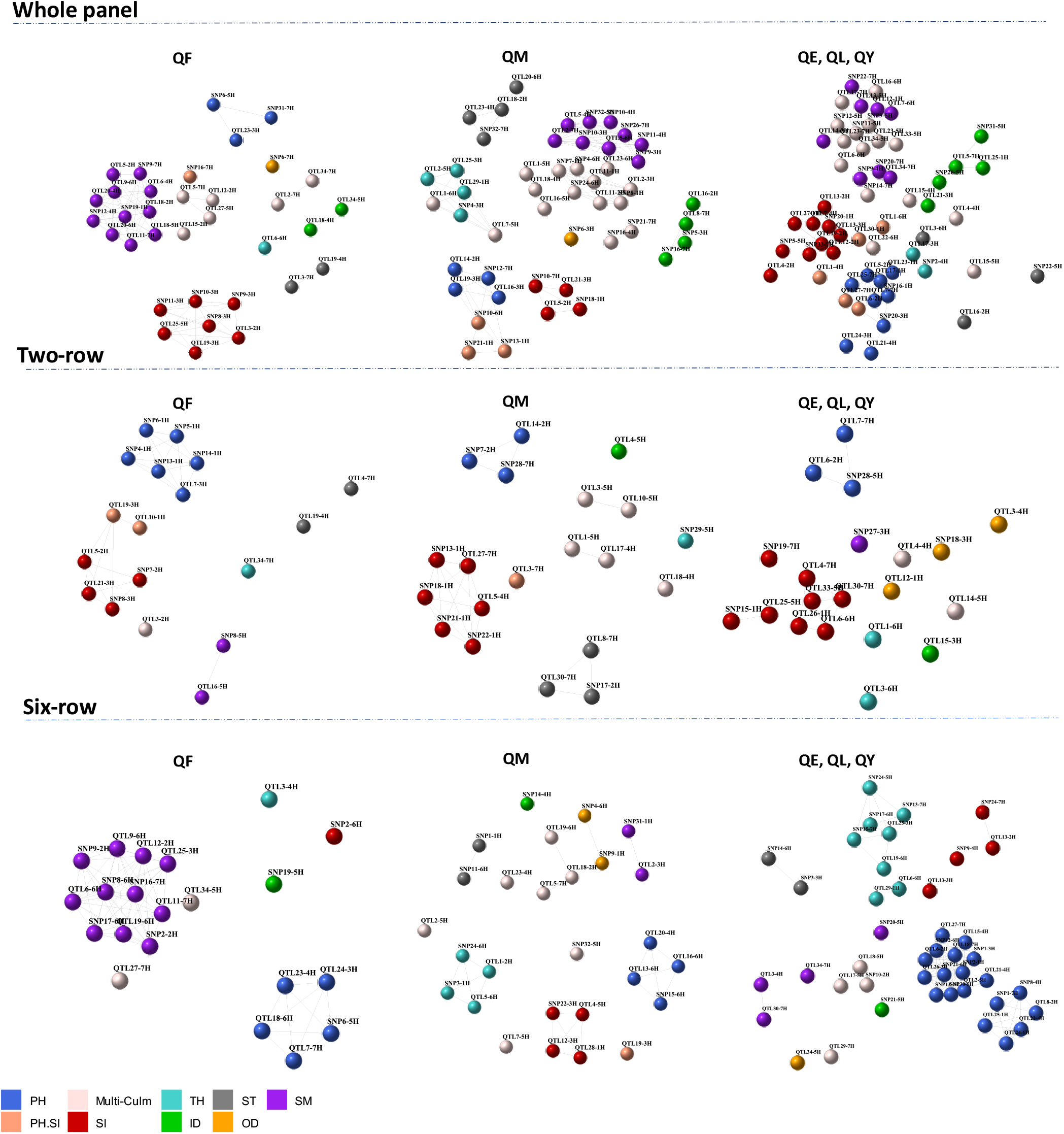
Co-association network representing co-association modules between 192 loci across whole panel and row type subsets, with color schemes according to the phenotypic traits. Each node is a SNP/QTL and a color according to its association with corresponding trait. Strong co-associations with a correlation above threshold (r = 0.9) are connected by edges.

Collectively, multi-environment GWAS results identified loci controlling culm morphology independent of plant height, with some QTLs showing stable effects across environments.

### Identification of QTLs with main and full effects and putative candidate gene exploration

In Table 2, we listed the most significant QTLs associated with the studied traits with QM or QF effects and potential candidate genes. The list of all 192 QTLs with complete details can be found in Supplemental Table S6 and synthetic view of genomic positions of QTLs along with the circular heatmap can be found in Figure 4 and Supplemental Figure S7. Promising candidate genes were selected based on literature searches, after excluding hypothetical genes and transposable elements. Marker-trait associations were listed with progressive numbering along chromosomes: as an example of the 93 loci detected by single SNPs, SNP1-1H is the first associated locus on chromosome 1H. The 99 QTLs detected by SNP clusters are designated as QTLs, e.g. QTL10-1H.

**Table 2.** Summary of candidate genes underlying the most significant markers with QM and QF effect on studied traits using multi-environment GWAS.

| QTL ID | Peak marker | Chr | Pos | QTL region (bp) | Panel | QTL type | Trait (-Log<sub>10</sub> P) | Gene | GeneID_MOREX.V2 |
| --- | --- | --- | --- | --- | --- | --- | --- | --- | --- |
| SNP4-1H | JHI-Hv50k-2016-19014 | 1H | 55564074 | 55564074 | 2-row | QF | PH (5.61) | *Asparticproteinase* | HORVU.MOREX.r2.1HG0014120 |
| SNP5-1H | JHI-Hv50k-2016-19267 | 1H | 59577895 | 59577895 | 2-row | QF | PH (5.61) | *Y14a* | HORVU.MOREX.r2.1HG0014550 |
| SNP7-1H | JHI-Hv50k-2016-21372 | 1H | 262131347 | 262131347 | Whole-panel | QM | OD (4.71), SM (5.66) | *HvCesA4/HvClsF4* | HORVU.MOREX.r2.1HG0032090 |
| QTL11-1H | JHI-Hv50k-2016-25138 | 1H | 331015882 | 331015882-332043281 | Whole-panel | QM | OD (4.71), SM (5.66) |  | HORVU.MOREX.r2.1HG0038730 |
| SNP13-1H | JHI-Hv50k-2016-26359 | 1H | 342720997 | 342720997 | 2-row | QF | PH (4.04) | *Rhmbd* | HORVU.MOREX.r2.1HG0039970 |
|  | JHI-Hv50k-2016-26359 | 1H | 342720997 |  | 2-row | QM | SI (5.49) |  |  |
|  | JHI-Hv50k-2016-26359 | 1H | 342720997 |  | Whole-panel | QM | PH (4.6), SI (5.08) |  |  |
| SNP18-1H | SCRI_RS_145336 | 1H | 423204614 | 423204614 | 2-row | QM | SI (5.72) | *UGT707A3* | HORVU.MOREX.r2.1HG0050860 |
|  | SCRI_RS_145336 | 1H | 423204614 |  | Whole-panel | QM | SI (5.96) |  |  |
| QTL29-1H | JHI-Hv50k-2016-52276 | 1H | 506206295 | 505927936-506431159 | Whole-panel | QM | TH (4.94) |  | Several candidate genes |
| QTL1-2H | 12_31446 | 2H | 1739468 | 1514860-1739468 | 6-row | QM | TH (5.01) | *SDG725* | HORVU.MOREX.r2.2HG0079410 |
| QTL5-2H | 12_31284 | 2H | 18622308 | 18522424-20868342 | 2-row | QF | SI (5.29) | *Eligulum-a* | HORVU.MOREX.r2.2HG0086910 |
|  | JHI-Hv50k-2016-71066 | 2H | 18623653 |  | Whole-panel | QM | SI (4.2) |  |  |
|  | JHI-Hv50k-2016-71911 | 2H | 20459215 |  | Whole-panel | QF | SM (5.66) |  |  |
| SNP7-2H | JHI-Hv50k-2016-75227 | 2H | 28446036 | 28446036 | 2-row | QF | SI (4.75) | *BRCT* | HORVU.MOREX.r2.2HG0090080 |
|  | JHI-Hv50k-2016-75227 | 2H | 28446036 |  | 2-row | QM | PH (4.99) |  |  |
| QTL11-2H | JHI-Hv50k-2016-103173 | 2H | 559241964 | 559231441-561722120 | Whole-panel | QM | SM (7.23) |  | Several candidate genes |
|  | JHI-Hv50k-2016-103187 | 2H | 559376614 |  | Whole-panel | QM | ID (5.7), OD (6.29) |  |  |
| QTL12-2H | JHI-Hv50k-2016-109824 | 2H | 598521913 | 597215106-598522113 | 6-row | QF | SM (4.27) |  | Several candidate genes |
|  | JHI-Hv50k-2016-109824 | 2H | 598521913 |  | Whole-panel | QF | ID (4.21), OD (4.8) |  |  |
|  | JHI-Hv50k-2016-109823 | 2H | 598522113 |  | Whole-panel | QF | SM (6.16) |  |  |
| QTL14-2H | JHI-Hv50k-2016-124833 | 2H | 633916743 | 633545278-635684841 | 2-row | QM | PH (4.55) |  | Several candidate genes |
|  | JHI-Hv50k-2016-124833 | 2H | 633916743 |  | Whole-panel | QM | PH (5.85) |  |  |
| QTL15-2H | JHI-Hv50k-2016-127337 | 2H | 638383926 | 638383926-638729272 | Whole-panel | QF | SM (5.12), TH (4.61) |  | Several candidate genes |
| QTL18-2H | JHI-Hv50k-2016-142360 | 2H | 665679591 | 665678929-669091745 | 6-row | QM | ID (5.09) |  | Several candidate genes |
|  | JHI-Hv50k-2016-142412 | 2H | 665806846 |  | Whole-panel | QM | ST (4.35) |  |  |
|  | JHI-Hv50k-2016-142417 | 2H | 665806970 |  | Whole-panel | QF | SM (4.18) |  |  |
|  | JHI-Hv50k-2016-142979 | 2H | 667239912 |  | 6-row | QM | SM (4.71) |  |  |

**Table 2.** Continued.
| QTL ID | Peak marker | Chr | Pos | QTL region (bp) | Panel | QTL type | Trait (-Log<sub>10</sub> P) | Gene | GeneID_MOREX.V2 |
| --- | --- | --- | --- | --- | --- | --- | --- | --- | --- |
| SNP9-3H | JHI-Hv50k-2016-167742 | 3H | 158248886 | 158248886 | Whole-panel | QM | SM (4.95) |  | - |
| QTL19-3H | JHI-Hv50k-2016-204951 | 3H | 570921209 | 570921209-571967329 | Whole-panel | QF | SI (4.26) | *HvGA20ox3* | HORVU.MOREX.r2.3HG0255700 |
|  | JHI-Hv50k-2016-204992 | 3H | 570930550 |  | 2-row | QF | PH (5.35), SI (4.99) |  |  |
|  | JHI-Hv50k-2016-205354 | 3H | 571967329 |  | 6-row | QM | PH (5.51), SI (5.74) |  |  |
|  | JHI-Hv50k-2016-205354 | 3H | 571967329 |  | Whole-panel | QM | PH (4.94) |  |  |
| QTL21-3H | JHI-Hv50k-2016-206708 | 3H | 579170749 | 577209764-583819782 | Whole-panel | QM | SI (5.44) |  | Several candidate genes |
|  | JHI-Hv50k-2016-207617 | 3H | 583528139 |  | 2-row | QF | SI (5.47) |  |  |
| QTL17-4H | JHI-Hv50k-2016-262348 | 4H | 586245828 | 586245828-586286949 | 2-row | QM | ID (4.68), OD (4.77), SM (4.77) | *CCD8d* | HORVU.MOREX.r2.4HG0337890 |
| QTL18-4H | JHI-Hv50k-2016-263046 | 4H | 589723289 | 589666126-590513139 | Whole-panel | QM | SM (4.95) |  | Several candidate genes |
|  | JHI-Hv50k-2016-263069 | 4H | 590012027 |  | 2-row | QM | SM (4.81) |  | - |
|  | JHI-Hv50k-2016-263069 | 4H | 590012027 |  | Whole-panel | QM | OD (5.05) |  |  |
|  | JHI-Hv50k-2016-263064 | 4H | 590012403 |  | 2-row | QM | TH (5.16) |  |  |
|  | JHI-Hv50k-2016-263064 | 4H | 590012403 |  | Whole-panel | QM | TH (4.73) |  |  |
|  | JHI-Hv50k-2016-263080 | 4H | 590144147 |  | 2-row | QM | OD (4.92) |  |  |
|  | JHI-Hv50k-2016-263116 | 4H | 590513139 |  | Whole-panel | QF | ID (4.07) |  |  |
| QTL19-4H | JHI-Hv50k-2016-263583 | 4H | 594808440 | 594808131-595015320 | Whole-panel | QF | ST (4.76) |  | Several candidate genes |
|  | JHI-Hv50k-2016-263787 | 4H | 594902360 |  | 2-row | QF | ST (4.86) |  |  |
| QTL23-4H | JHI-Hv50k-2016-275313 | 4H | 621344288 | 621344288-622035884 | 6-row | QM | ID (5.33), ST (4.75) |  | Several candidate genes |
|  | JHI-Hv50k-2016-275693 | 4H | 621902266 |  | 6-row | QF | PH (4.71) |  |  |
|  | JHI-Hv50k-2016-275696 | 4H | 621902455 |  | Whole-panel | QM | ST (4.16) |  |  |
| QTL1-5H | JHI-Hv50k-2016-277297 | 5H | 1444564 | 869533-2211050 | Whole-panel | QM | OD (4.26), SM (4.2) | *CCD1* | HORVU.MOREX.r2.5HG0349440 |
|  | JHI-Hv50k-2016-277332 | 5H | 1447495 |  | Whole-panel | QM | TH (4.33) |  |  |
|  | JHI-Hv50k-2016-277338 | 5H | 1448582 |  | 2-row | QM | SI (4.09) |  |  |
|  | JHI-Hv50k-2016-277724 | 5H | 2211050 |  | 2-row | QM | ID (5.05), OD (5.6), SM (5.62) |  |  |
| QTL2-5H | JHI-Hv50k-2016-278616 | 5H | 4053376 | 3330549-5170277 | 6-row | QM | ST (5.33), TH (4.62) |  | Several candidate genes |
|  | JHI-Hv50k-2016-278616 | 5H | 4053376 |  | Whole-panel | QM | TH (4.58) |  |  |
| QTL3-5H | JHI-Hv50k-2016-279858 | 5H | 6140678 | 6139160-6687421 | 2-row | QM | ID (4.7) |  | Several candidate genes |
| QTL4-5H | JHI-Hv50k-2016-281676 | 5H | 10305211 | 10221340-10615460 | 6-row | QM | SI (5.24) |  | Several candidate genes |
|  | JHI-Hv50k-2016-281715 | 5H | 10326076 |  | 2-row | QM | ID (4.12) |  |  |

**Table 2.** Continued.
| QTL ID | Peak marker | Chr | Pos | QTL region (bp) | Panel | QTL type | Trait (-Log<sub>10</sub> P) | Gene | GeneID_MOREX.V2 |
| --- | --- | --- | --- | --- | --- | --- | --- | --- | --- |
| QTL7-5H | JHI-Hv50k-2016-287215 | 5H | 27977719 | 27977719-33923320 | 6-row | QM | SI (4.76) |  | Several candidate genes |
|  | JHI-Hv50k-2016-287531 | 5H | 29346346 |  | Whole-panel | QM | TH (4.12) |  |  |
|  | JHI-Hv50k-2016-287643 | 5H | 29357949 |  | Whole-panel | QM | SI (4.03) |  |  |
|  | JHI-Hv50k-2016-288619 | 5H | 33923127 |  | 6-row | QM | TH (5.47) |  |  |
| QTL16-5H | JHI-Hv50k-2016-309388 | 5H | 452787194 | 449553280-453741857 | Whole-panel | QM | TH (6.32) |  | Several candidate genes |
|  | JHI-Hv50k-2016-309383 | 5H | 452787694 |  | 2-row | QF | SM (4.3) |  |  |
|  | JHI-Hv50k-2016-309383 | 5H | 452787694 |  | Whole-panel | QM | OD (5.48), SI (4.19), SM (6.94) |  |  |
| SNP19-5H | JHI-Hv50k-2016-310560 | 5H | 467079429 | 467079429 | 6-row | QF | ID (5.79) | *ABA8ox2* | HORVU.MOREX.r2.5HG0402930 |
| QTL27-5H | JHI-Hv50k-2016-329041 | 5H | 525702571 | 525598290-525702571 | Whole-panel | QF | ID (5.78), OD (5.23), SM (6.00) | *SIP1* | HORVU.MOREX.r2.5HG0420210 |
| SNP32-5H | 12_31206 | 5H | 553957781 | 553957781 | 6-row | QM | OD (4.41), SI (7.62), SM (5.63) | *SAP6* | HORVU.MOREX.r2.5HG0430170 |
|  | 12_31206 | 5H | 553957781 |  | Whole-panel | QM | SM (4.05) |  |  |
| QTL6-6H | JHI-Hv50k-2016-383797 | 6H | 36026739 | 35725637-37076534 | Whole-panel | QF | TH (5.23) |  | - |
| SNP10-6H | SCRI_RS_161533 | 6H | 242933786 | 242933786 | Whole-panel | QM | PH (4.93), SI (4.22) | *UBP15, LG1* | HORVU.MOREX.r2.6HG0483350 |
| QTL13-6H | JHI-Hv50k-2016-405999 | 6H | 431754022 | 428846608-435119247 | 6-row | QM | PH (5.18) |  | Several candidate genes |
| SNP17-6H | 12_30573 | 6H | 512709462 | 512709462 | 6-row | QF | SM (5.66) | *RFP* | HORVU.MOREX.r2.6HG0509750 |
| QTL3-7H | JHI-Hv50k-2016-449409 | 7H | 13356822 | 12920299-14593868 | 2-row | QM | SI (5.25) | *SMOS2/DLT* | HORVU.MOREX.r2.7HG0534100 |
|  | JHI-Hv50k-2016-449409 | 7H | 13356822 |  | Whole-panel | QF | ST (5.87) |  |  |
|  | JHI-Hv50k-2016-449626 | 7H | 13692220 |  | 2-row | QM | ST (5.95) |  |  |
| QTL5-7H | JHI-Hv50k-2016-453012 | 7H | 22070216 | 21643770-22444585 | Whole-panel | QF | OD (4.99), SM (5.03) |  | Several candidate genes |
|  | JHI-Hv50k-2016-453082 | 7H | 22441304 |  | 6-row | QM | ID (4.7), OD (4.52), SM (4.12) |  |  |
| QTL7-7H | JHI-Hv50k-2016-460460 | 7H | 39722386 | 38675923-39722386 | 6-row | QF | PH (4.82) | *HvFT1/VRNH3* | HORVU.MOREX.r2.7HG0542540 |
| SNP16-7H | JHI-Hv50k-2016-478948 | 7H | 265292093 | 265292093 | 6-row | QF | SM (4.18) | *NTL* | HORVU.MOREX.r2.7HG0573190 |
|  | JHI-Hv50k-2016-478948 | 7H | 265292093 |  | Whole-panel | QF | OD (5.97), PH (4.46), SM (7.28) |  |  |
|  | JHI-Hv50k-2016-478948 | 7H | 265292093 |  | Whole-panel | QM | ID (6.33) |  |  |
| QTL27-7H | SCRI_RS_168994 | 7H | 570828407 | 570827595-572601830 | 6-row | QF | OD (4.74), SM (5.72) | *DWARF27* | HORVU.MOREX.r2.7HG0603370 |
|  | JHI-Hv50k-2016-493265 | 7H | 572601830 |  | 2-row | QM | SI (4.54) |  |  |
| QTL30-7H | JHI-Hv50k-2016-501203 | 7H | 598638988 | 597448728-600244977 | 2-row | QM | ST (4.94) | *DEP3* | HORVU.MOREX.r2.7HG0610260 |
| SNP32-7H | SCRI_RS_213791 | 7H | 625219043 | 625219043 | Whole-panel | QM | ST (5.05) |  | HORVU.MOREX.r2.7HG0620190 |

**Table 2.** Continued.
| QTL ID | Peak marker | Chr | Pos | QTL region (bp) | Panel | QTL type | Trait (-Log<sub>10</sub> P) | Gene | GeneID_MOREX.V2 |
| --- | --- | --- | --- | --- | --- | --- | --- | --- | --- |
| QTL34-7H | JHI-Hv50k-2016-516642 | 7H | 628347284 | 628346780-633832080 | Whole-panel | QF | ID (4.27), OD (4.07) | *HvDIM* | HORVU.MOREX.r2.7HG0622270 |
|  | JHI-Hv50k-2016-518794 | 7H | 632545446 |  | Whole-panel | QF | TH (5.15) |  |  |
|  | JHI-Hv50k-2016-519440 | 7H | 633832080 |  | 2-row | QF | TH (5.49) |  |  |

**Figure 4.**
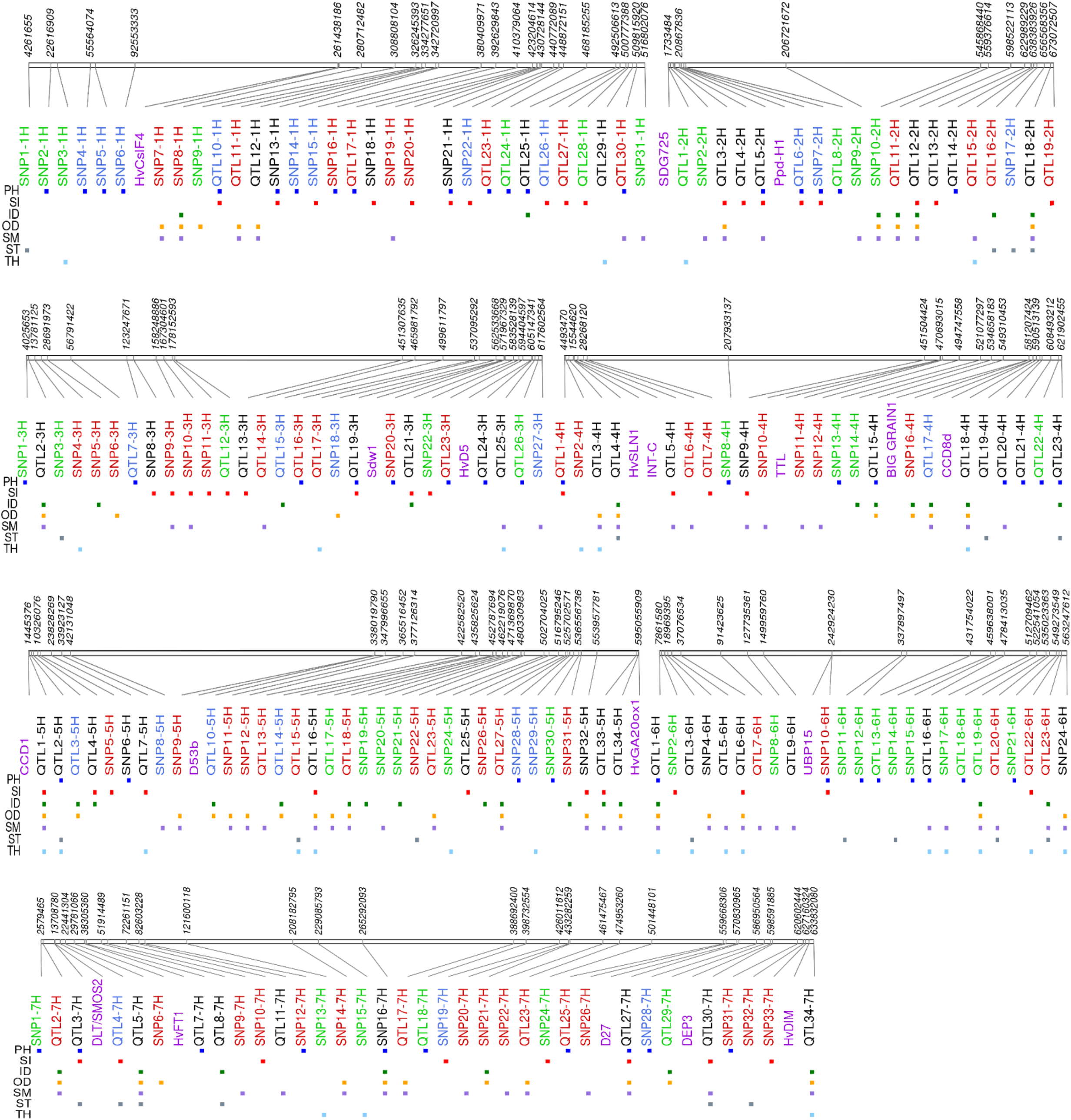
Physical map of 192 QTLs associated with culm morphological traits a cross whole panel and the row type subsets. Loci with red, blue, and green colors are unique to whole panel, two-row, and six-row subsets, respectively. Loci with black color are those detected at least in two association panel. Purple color indicates relative position of barley known genes at that particular genomic region.

Out of a total of 31 QTLs on chromosome 1H, the most significant were SNP4-1H, SNP5-1H, SNP7-1H, SNP8-1H, and QTL11-1H. SNP7-1H (pos: 262.13 Mb) was associated with both OD and SM in the whole panel and located in close proximity with candidate gene *HvCesA4/HvClsF4*, encoding a cellulose synthase protein previously associated to culm strength in barley (Burton et al., 2010).

For chromosome 2H, 19 QTLs were detected. QTL1-2H associated with TH (six-row panel) explained a high proportion of phenotypic variance. We found that QTL1-2H (pos: 1.51 – 1.74 Mb) harbors the ortholog of rice *OsSDG725* encoding a histone H3K36 methyltransferase and playing an important role in rice plant growth and development (Sui et al., 2012).

For chromosome 3H, 27 QTLs were identified including QTL19-3H (pos: 570.92 - 571.97 Mb) which is associated with both PH and SI across all panels and spans the well-known plant height gene *Sdw1* (Xu et al., 2017) found in many elite European 2-row spring barley cultivars.

On chromosome 4H, a total of 23 QTLs were identified. A particularly interesting region with QM effect was QTL17-4H (pos: 586.24 -586.29 Mb) associated with ID, OD, and SM in the two-row panel and explaining at least 6% of the phenotypic variance. This QTL was found to harbor a homolog of rice *CCD8-d* (carotenoid cleavage dioxygenase). QTL18-4H was detected in both the whole panel (ID, OD, SM, TH) and two-row panel (ID, OD, SM), explaining between 2.74% to 7.1% of variance (pos: 589.66 Mb – 590.52 Mb). SNP10-4H was associated with SM and located within a pseudo-response regulator gene (470.68 Mb). Also, about 0.8 Mb from this marker we noted a homolog of *TRANSTHYRETIN-LIKE PROTEIN* (*TTL*), a gene that was previously associated with stem circumference in sorghum (Mantilla Perez et al., 2014). OD and ID were associated with SNP16-4H (481.27 Mb), 0.5 Mb from a homolog of rice *BIG GRAIN1* (Liu et al., 2015).

On chromosome 5H, 34 QTLs were detected, including three loci with promising associations. QTL1-5H was identified in two-row panel as associated with ID, OD, SM, SI (pos: 0.87 - 2.21 Mb) and contained the rice homolog of *OsCCD1* (Ilg et al., 2009). QTL2-5H predominantly associated in the six-row panel with PH, ST, TH and in the whole panel for TH (pos: 3.33 – 5.17 Mb) and explained more than 8% of variance for TH and ST in the six-row panel and harbors several uncharacterized genes. SNP32-5H (pos: 553.95 Mb) was associated with OD, SI, and SM in both the six-row and the whole panel.

For chromosome 6H, in total 24 QTLs were identified, among them there were two SNPs with promising effect. SNP10-6H associated with both PH and SI at position 242.933 Mb located within a gene encoding a ubiquitin carboxyl-terminal hydrolase closely related to rice *Large Grain 1* (*LG1*/*OsUBP15*), a gene involved in seed size and plant height (Shi et al., 2019). SNP17-6H (512.71 Mb) was associated with SM and TH and falls within an uncharacterized gene encoding a RING/U-box superfamily protein. A large QTL region, QTL13-6H, was associated with PH in the six-row panel (pos: 428.84 – 435.12 Mb) and contains several uncharacterized genes.

On chromosome 7H, a total of 34 QTLs were detected including six QTLs of special interest. QTL3-7H, was associated with ST in the whole panel, and PH, SI, and ST in the two-row panel (pos: 12.92 Mb – 14.59 Mb). The region contains several candidates including a gene encoding a GRAS transcription factor orthologous to rice *DWARF AND LOW-TILLERING* (*DLT*/*SMOS2*), that can directly interact with *SMALL ORGAN SIZE1* (*SMOS1*/*RLA1*), and *RLA1* plays as an integrator with both *OsBZR1* and *DLT* to modulate their activity (Tong et al., 2009; Tong et al., 2012; Hirano et al., 2017b; Qiao et al., 2017).

QTL5-7H was associated with ID, OD, and SM in both whole and six-rows panels and also with ST in the six-row panel (pos: 21.64 – 22.45 Mb). SNP16-7H (pos: 265.29 Mb), a hotspot SNP associated with ID, OD and SM in the six-row and whole panels. Another noteworthy QTL was QTL27-7H, associated with PH, SI in the whole panel, OD, PH and SM in the six-row panel, and SI in the two-row panel (pos: 570.827 Mb – 572.61 Mb). The region contains *HvD27*, the barley ortholog to rice strigolactone biosynthesis gene *DWARF27* encoding beta-carotene isomerase (Lin et al., 2009). QTL30-7H (pos: 597.44 Mb – 600.25 Mb) was associated with SI, SM, and TH and contains several genes including a patatin encoding protein gene highly related to *DEP3*, a rice gene previously shown to affect culm morphology and anatomy as well as panicle architecture (Qiao et al., 2011). Finally, QTL34-7H (pos: 628.34 Mb – 633.84 Mb) was associated with TH in the two-row panel and with ID, OD, and TH in the whole panel. This locus had also QL effect with SM both in the six-row and whole panel and contains *HvDIM* encoding Delta(24)-sterol reductase previously shown to act in the brassinosteroid pathway in barley (Dockter et al., 2014)).

### Identification of QTLs with interaction effects

Besides the above-mentioned QTLs with main and full effects, multi-environment GWAS uncovered highly significant QTLs with interaction effects. QTL26-1H (pos: 495.79 – 497.02 Mb) was associated with SI in the two-row panel. QTL6-2H (pos: 22.37 – 23.99 Mb) associated with SI and PH (whole, two-row, and six-row panels) spans the well-known barley *PPD-H1* gene (Supplemental Table S6), involved in photoperiod responsive flowering (Turner et al., 2005). The genomic region of QTL15-3H (pos: 499.61 – 499.87 Mb), associated with ID in two-row subset, hosted uncharacterized genes. QTL34-5H (pos: 594.17 – 596.71 Mb) was associated with ID, OD, and SM in the whole and six-row panels. This QTL showed QE and QF effects in the whole panel and six-row panel, respectively, and contains a barley Gibberellin 20 oxidase, *HvGA20ox1*, which has recently been associated to straw breaking and flowering time in barley (Göransson et al., 2019; He et al., 2019). QTL7-7H for PH was found across all panels and located in close proximity to the barley *HvFT1*/*VRNH3* gene. It showed QL effect in the whole and two-row panels and QF effect in the six-row panel. In barley, *HvFT1* expression requires the active version of *PPD-H1* to promote flowering under long day conditions (Hemming et al., 2008). Currently there is no report on its effect on plant height.

### Allelic comparison of SNPs/QTLs with QM/QF effects for lodging and plant height

In order to appraise the effects of the QTLs on lodging susceptibility, we focused on QTLs with QM and QF effects (Supplemental Figures S8 and S9, respectively, Supplementary Table S6). Allelic comparisons for these loci indicated that depending on the trait and sub-population their effect was highly variable. As expected, QTLs for PH and SI showed significant differences for both PH and LG. With respect to culm morphology QTLs, effects on PH and LG were variable ranging from no difference to significant differences, including some QTLs that significantly affected both LG and PH. However, most QTLs associated with culm morphology had no effects on PH in the whole panel, but showed significant effects on LG. Such types of QTLs were also detected in both six-row and two-row panels. For example, the QTLs associated with ID, OD, and SM -SNP8-1H, QTL11-1H, QTL11-2H, QTL2-3H, SNP16-4H, QTL16-5H, QTL5-6H, QTL26-6H, QTL2-7H, SNP26-7H-affected lodging without any effect on PH in the whole panel. In two-row, some examples are QTL17-4H, QTL18-4H, QTL10-5H, and QTL16-5H. Finally, for six-row panel SNP9-1H, SNP14-4H are among the QTLs affecting lodging without any effect on PH. Considering loci with main effects (Supplemental Figure S8), out of 20 loci associated to OD, 11 had a significant impact on LG without any effect on PH (8 in the whole panel, 1 and 2 in the six-row and two-row, respectively), and out of 25 loci detected for SM, 16 significantly affected LG without impacting PH (14 in the whole panel, 2 in two-row): nine of these QTLs were shared between OD and SM (SNP7-1H, SNP8-1H, QTL11-1H, QTL11-2H, QTL2-3H, QTL17-4H, QTL18-4H, QTL16-5H, SNP32-5H). Interestingly, QTL18-4H was detected in both the whole panel and the two-row panel also for TH, indicating this locus as an interesting target for manipulation of culm morphology and lodging resistance. However, fewer loci associated with TH and ST had effects on LG. We thus focused on OD, ID and SM for more detailed analyses of nine SNPs associated with these traits in the whole panel: SNP7-1H, SNP8-1H, SPN5-3H, SNP10-4H, SNP11-4H, SNP16-4H, SNP32-5H, SNP21-7H and SNP26-7H. In all cases, alleles increasing culm diameter (OD, ID) and/or SM had negative effects on lodging, without affecting PH (Table 3).

**Table 3.** Details of subset of SNPs with main effects and associated with culm traits with negative effects on lodging without impacting on plant height. A1, A2, and MAF indicate major allele, minor allele and minor allele frequency, respectively. The allele associated with decreased lodging is underlined. PVE (%) is the percentage of phenotype variance explained by SNP. β is the SNP main effect, α_1_ is the SNP -by-location effect, and α_2_ is the SNP -by-year effect derived from GWAS model.

| SNP | Panel | Trait | Peak marker | MAF | A1 | A2 | chr | position | -Log10<br>(P-value) | $\beta$ | $\alpha_1$ | $\alpha_2$ | PVE<br>(%) |
| --- | --- | --- | --- | --- | --- | --- | --- | --- | --- | --- | --- | --- | --- |
| SNP7-1H | Whole panel | OD | JHI-Hv50k-2016-21372 | 0.13 | A | <u>C</u> | 1H | 262131347 | 4.71 | 0.472 | -0.012 | -0.001 | 2.46 |
|  |  | SM |  | 0.13 | A | <u>C</u> | 1H | 262131347 | 5.66 | 0.533 | -0.024 | 0.008 | 3.41 |
| SNP8-1H | Whole panel | ID | JHI-Hv50k-2016-22255 | 0.14 | C | <u>A</u> | 1H | 280712482 | 4.15 | 0.481 | -0.008 | -0.012 | 2.91 |
|  |  | OD |  | 0.14 | C | <u>A</u> | 1H | 280712482 | 4.26 | 0.502 | -0.01 | 0.007 | 2.98 |
|  |  | SM |  | 0.14 | C | <u>A</u> | 1H | 280712482 | 4.15 | 0.501 | -0.019 | 0.022 | 3.24 |
| SNP5-3H | Whole panel | ID | JHI-Hv50k-2016-162361 | 0.21 | <u>A</u> | G | 3H | 28691973 | 4.06 | -0.255 | 0.003 | 0.003 | 1.53 |
| SNP10-4H | Whole panel | SM | JHI-Hv50k-2016-246906 | 0.09 | C | <u>T</u> | 4H | 470693015 | 4.21 | 0.554 | -0.035 | 0.02 | 1.72 |
| SNP11-4H | Whole panel | SM | JHI-Hv50k-2016-247273 | 0.09 | G | <u>T</u> | 4H | 474202180 | 4.21 | 0.554 | -0.035 | 0.02 | 1.72 |
| SNP16-4H | Whole panel | ID | JHI-Hv50k-2016-261211 | 0.15 | T | <u>C</u> | 4H | 581266705 | 4.38 | 0.307 | -0.007 | 0.003 | 1.23 |
|  |  | OD |  | 0.15 | T | <u>C</u> | 4H | 581266705 | 4.30 | 0.312 | 0.001 | 0.002 | 1.21 |
| SNP32-5H | Whole panel | SM | 12_31206 | 0.27 | C | <u>G</u> | 5H | 553957781 | 4.05 | 0.185 | 0 | -0.005 | 1.06 |
|  | Six-row | OD |  | 0.24 | C | <u>G</u> | 5H | 553957781 | 4.41 | 0.345 | -0.005 | 0.002 | 3.51 |
|  | Six-row | SI |  | 0.24 | C | <u>G</u> | 5H | 553957781 | 7.62 | 0.522 | 0.017 | -0.042 | 6.62 |
|  | Six-row | SM |  | 0.24 | C | <u>G</u> | 5H | 553957781 | 5.63 | 0.358 | 0 | -0.006 | 4.85 |
| SNP21-7H | Whole panel | ID | JHI-Hv50k-2016-486762 | 0.19 | <u>C</u> | G | 7H | 434555860 | 4.26 | -0.482 | -0.011 | -0.005 | 4.93 |
|  |  | OD |  | 0.19 | <u>C</u> | G | 7H | 434555860 | 4.05 | -0.481 | -0.004 | -0.02 | 4.48 |
| SNP26-7H | Whole panel | SM | JHI-Hv50k-2016-492337 | 0.24 | <u>C</u> | T | 7H | 562028351 | 4.23 | -0.511 | 0.017 | -0.034 | 7.02 |

In conclusion, results from these analyses support the usefulness of SM and culm diameter as parameters for selecting alleles to improve lodging resistance and provide chromosomal positions and markers associated to promising loci.

## Discussion

In the present study, we investigated natural genetic variation for morphological characteristics of the barley culm and their relationships with lodging and agronomic traits. To date, no genetic studies have used image-based phenotyping to investigate the genetic architecture of culm morphology in barley. For this reason, we developed a robust method to extract quantitative measurements of culm diameter and thickness from images of culm sections, showing that significant phenotypic variation exists within our barley germplasm panel with a major contribution of genetic variation to these traits as supported by medium-high heritability values.

Using PCA we showed that row-type and germplasm source are the major factors driving population structure of the panel. In addition, no evidence of strong admixture between row-type groups was observed in PCA. This is consistent with previous studies suggesting that breeders largely focused within the six-row and two-row gene pools in developing new varieties therefore limiting the exchange of genetic variation between these major row-types, despite some cases of targeted introgression (Hernandez et al., 2020). Increasing seed number per spike was probably the reason for the human selection of recessive allele at *VRS1* into the barley gene pool during domestication (Komatsuda et al., 2007). On the other hand, barleys most commonly grown in Europe are two-row cultivars, which are preferred for malting because of uniformity in seed size: this resulted in limited genetic diversity compared to the six-row cultivars. This variation in seed size is due mainly to the allelic variation at the *INT-C/VRS5* gene between row-types (Ramsay et al., 2011). Row-type genes have pleiotropic effects on other traits, as well-known for tillering (Liller et al., 2015). In our study, row-type subsets exhibited clear differences also for some culm morphological traits, e.g. six-row barleys showed higher mean values of TH and SM compared to two-row barleys. Relationships between row-type and the studied traits are also evident from positive correlations with PH, OD, ID, SM, TH, LG and negative correlations with HD, ST, SI.

Correlation results showed that although plant height is important for lodging, culm characteristics also play an important role in lodging resistance. We observed strong positive correlations among culm traits, as well as negative correlations between culm traits and lodging across the whole and row-type panels. On the other hand, culm morphological traits showed weak (two-row) and even no (six-row panel) correlations with plant height. This suggests opportunities for genetic improvement of lodging resistance through manipulation of culm morphology independent of plant height. Generally, relationships among traits were similar across row-type subpopulations, sometimes with different magnitudes: for example, correlation between LG and OD was -0.52 and -0.69 in two- and six-row subpanels, respectively. Interesting correlations specific of the six-row subset were detected between LG, PH and HD with six-row landraces being late heading, taller and more prone to lodging compared to six-row cultivars: these landraces also had lower values of OD, ID and SM, therefore combining different unfavorable traits for lodging susceptibility. It should be noted that these contrasting patterns may be due to the fact that the six-row cultivars were mainly early flowering lines of Scandinavian origin, while the six-row landraces were of Mediterranean origin. Based on these observations, it would be interesting to further explore the genetic relationships between heading, plant height and culm morphological traits in a wider sample of six-row barleys.

Based on these results, we analyzed phenotypic variation and run mixed model GWAS in the whole panel, as well as row-type subgroups independently in order to: i) minimize the confounding effects of row-type on association analyses; ii) understand if distinct loci are segregating in row-type subpopulations and thus different regulatory networks are involved in genetic control of the studied traits. The use of mixed model in GWAS is a well-established approach to efficiently reduce false positive associations for most traits, but it may also mask true signals that are correlated with population structure. As a result, loci that distinguish barley subpopulations are often difficult to detect using mixed model. To circumvent this problem, many association mapping studies have analyzed each subpopulation separately and successfully identified loci specific to each subpopulation. In our study, 120 marker-trait associations were detected in the whole panel, including 21 and 27 that were shared with the two-row and six-row panels respectively. Six associations were detected across all three panels. In addition, we uncovered 24 and 45 QTLs specific for two- and six-row panels, supporting the relevance of running GWAS on row-type subsets. We also noticed that for some QTLs detected across both row-types, allele frequencies and peak markers differed between the row-type subsets, resulting in opposite effects of minor alleles on the same trait. Taking as an example the PH locus QTL19-3H spanning the well-characterized *Sdw1* gene, the peak marker in the six-row panel was JHI-Hv50k-2016-205354 with the minor allele showing a negative effect on PH and positive effect on SI, in contrast to the effect of JHI-Hv50k-2016-204992, the peak marker in the two-row panel. Likewise, QTL6-2H containing *PPD-H1* had negative effects on PH in the six-row panel while the effect in two-row was the opposite. This indicates that causative variants in these major genes have different frequencies and are associated to different markers in row-type subsets. Taken together, comparative analysis of results from the whole panel and row-type subsets indicates the need to duly account for population structure in dissecting culm morphological traits and carefully analyze effects of potentially interesting markers for breeding in relation to row-type population. This is also relevant when considering crosses between row types in the context of plant breeding.

While in our GWAS analysis we observed numerous trait-specific QTLs, we also observed QTLs that were associated with multiple traits. In addition, in the same QTL region, the peak marker was sometimes different depending on the panel. For example, QTL34-7H was associated with OD, ID, SM, TH in the whole panel, with SM in the six-row panel, and with TH in two-row panel (QF effect, Supplemental Figure S8). The peak markers for TH were also different between the whole panel and the two-row panel, while the peak marker for SM was common between six-row and the whole panel. This QTL harbors the *HvDIM* gene encoding the barley Δ5-sterol-Δ24-reductase, an enzyme involved in the brassinosteroid biosynthetic pathway (Dockter et al., 2014). A link between brassinosteroids and culm thickness is supported by studies of the rice *SMOS1* and *SMOS2* genes, encoding transcription factors of the AP2 and a GRAS family, respectively, that interact to integrate auxin and brassinosteroid signaling: *smos1* and *smos2* single mutants as well as *smos1-smos2* double mutants show increased culm thickness (Hirano et al., 2017b). Classical semidwarf barley mutants *brh*.*af, brh14*.*q, brh16*.*v, ert-u*.*56, ert-zd*, and *ari*.*o* were shown to harbor mutations in the *HvDIM* gene (Dockter et al., 2014): these mutants have reduced plant height and are more resistant to lodging compared to respective wild type (Dahleen et al., 2005), but their culm morphological traits have not been described. In our work a marker within this region showed weak association with PH (JHI-Hv50k-2016-516979, p value=0.003), suggesting *HvDIM* as a possible candidate for QTL34-7H. However, there are other potential candidates in this genomic region that have been reported as members of glycosyl transferase (GT) gene family, such as cellulose synthase genes of the GT2 family that influences culm cellulose content (Houston et al., 2015). Given the significance of associations between this genomic region and multiple culm morphology traits, it would be interesting to further dissect this QTL to discriminate if such effects are the result of pleiotropy or closely linked genes (local LD) and identify the underlying gene(s)/alleles combining association mapping and biparental fine mapping.

Taking advantage of data from seven different environments, multi-environment GWAS (Korte et al., 2012) enabled us to disentangle QTLs with main effects stable across environments (QM) from QTLs with environment-dependent effects (location and/or year). An example of a QTL with a significant interaction with location is QTL6-2H, which was detected for PH across all panels. This genomic region contains the well-known *PPD-H1* gene (Turner et al., 2005), a major regulator of barley flowering in response to photoperiod, that was shown to have pleiotropic effects on several agronomic traits including yield, leaf size and plant height (Karsai et al., 1999; Digel et al., 2016). With respect to lodging, alleles with stable phenotypic effects across environments are preferable for breeding under changing climatic conditions. For this reason, we decided to focus our attention on culm morphology QTLs with main effects, showing significant negative impact on lodging without affecting PH: for nine SNPs detected in the whole panel, alleles increasing culm diameter and/or SM consistently reduced lodging (SNP7-1H, SNP8-1H, SNP5-3H, SNP10-4H, SNP11-4H, SNP16-4H, SNP32-5H, SNP21-7H and SNP26-7H). We scanned regions adjacent to these SNPs ± 0.8 Mb (i.e. the genome-wide LD decay estimated for the whole panel) in order to search for potential candidate genes. For example, cellulose synthase gene *HvCslF4* (1H, 261.4 Mb) is located near SNP7-1H (262.1 Mb): a retroelement insertion within this gene was previously associated with the *fragile stem2* (*fs2*) mutant phenotype in barley, suggesting a link between stem strength and genes involved in cellulose content (Burton et al., 2010). Since we analyzed culm morphology traits in straw culm sections, cell wall composition and cellulose content are likely to impact the morphological features considered in our work. Another example is SNP32-5H (5H, 553.9 Mb): the adjacent region hosts several possible candidate genes, including *HvMND1* (552.9 Mb), which encodes a N-acetyl-transferase-like protein recently shown to regulate barley plastochron and plant architecture (Walla et al., 2020).

Beside these SNPs, additional QTLs were identified as associated with culm features and having impact on lodging, independent of PH. Among them, QTL17-4H had main effects on ID, OD and SM and contained a carotenoid cleavage dioxygenase 8 (*CCD8*) gene located in close proximity to the peak marker. A recent phylogenetic study showed that rice has four *CCD8* genes (*CCD8-a, -b, -c*, and *-d*), while Arabidopsis has only one: both Arabidopsis *CCD8* and rice *CCD8-b* are involved in the biosynthesis of strigolactones, phytohormones that control lateral shoot growth, and affect stem thickness at least in some species (reviewed in Chesterfield et al., 2020). The barley ortholog of *OsCCD8-b* is located on chromosome 3H, while the *CCD8* gene associated with QTL17-4H is more closely related with *OsCCD8d*, whose function has not been characterized yet (Priya and Siva, 2014). An alternative candidate gene for this QTL may be *MDP1*, encoding a MADS box transcription factor implicated in brassinosteroid signaling (Duan et al., 2006).

While validation of these potential candidate genes will require more detailed analyses, our results provide the first insights into the genetic architecture of culm morphology in barley and its relevance for lodging. Utilization of loci underpinning culm features may open new avenues to improve lodging resistance and increase barley yield stability under changing environments.

## Materials and methods

### Plant materials, experimental design and phenotyping

The germplasm collection considered in this study was composed of 165 two-row and 96 six-row barley lines, including both European cultivars and a set of Spanish landraces (Supplemental Table S1) grown at two Northern and two Southern European sites respectively. Southern sites were winter-sown and for these sites only we included 34 Spanish landraces that had a vernalisation requirement. Barley lines were sown for two consecutive harvest years, 2016 and 2017, in four European research stations (Supplemental Table S3), except for the LUKE site (Finland), where data were collected only for 2017. Fields were organized in row and column designs with 2 complete replicates. Each plot covered on average 2m^2^ and all the trials were rainfed - additional details about field trials and sowing densities are presented in Supplemental Table S3.

Zadoks scale was used throughout all trials in order to define the specific developmental stage for sampling and phenotypic measurements (Zadoks et al., 1974). Details of phenotyping methods used to measure the studied traits are described in Supplemental Table S4. Samples were collected from plot centres at Zadoks stage 90 from the second internode of the main culm, which is considered a critical area for lodging resistance (Pinthus, 1974; Berry et al., 2004). A dedicated image analysis-based protocol was developed for measurement of culm morphological traits and additional details can be found in Supplemental Methods S1.

### Genome-wide SNPs genotyping and genotype imputation

The barley germplasm panel was genotyped with the 50k Illumina Infinium iSelect genotyping array (Bayer et al., 2017). Physical positions of markers were based on pseudo-molecule assembly by Monat et al., 2019. Allele calls were made using GenomeStudio Genotyping Module v2.0.2 (Illumina, San Diego, California). After manual checking, SNP markers with more than two alleles, missing values greater than 10%, minor allele frequency (MAF) < 5% were excluded from analyses, along with unmapped SNPs. As a result, 36020 SNP markers and 261 genotypes (165 two-row and 96 six-row barleys) remained for the analysis. Missing genotypes were imputed using Beagle v5.0 (Browning et al., 2018, Supplemental Methods S2).

### Linkage disequilibrium, population structure, and kinship

LD is in many cases influenced by the presence of population structure and relatedness due to demographic and breeding history of the accessions. To take into consideration these factors, the intrachromosomal LD between two SNPs was estimated as squared allele-frequency correlations (r^2^) using an unbiased (due to non-independence relationships between individuals) estimation implemented in the R package called LDcorSV (Mangin et al., 2012). The markers were thinned to every three SNP and LD between all pairs of intrachromosomal sites was estimated. Four r^2^ estimates were calculated: r^2^ based on raw genotype data, r^2^ with population structure represented by the PCA after scaling the PC scores across a range of zero to one (rs2, see below), r^2^ with relatedness (rv2; see the next section), and r^2^ with both population structure and relatedness (rsv2). The r^2^ values were plotted against the physical distance (Mb) and a nonlinear LOESS curve was fitted to investigate the relationship between LD and physical distance. A square root transformation of unlinked r^2^ values was calculated and the parametric 95th percentile of the distribution of transformed values was taken as a critical r^2^ value (Breseghello and Sorrells, 2006). The unlinked r^2^ refers to the r^2^ between the SNP loci with a physical distance greater than 50 Mb.

Population structure was estimated using principal components analysis. Prior to PCA, the genotype marker data were filtered out by LD-pruning to generate a pruned dataset of SNPs that are in approximate linkage equilibrium, thus reducing the effect of LD on population structure. The LD-based SNP pruning was conducted with a window size of 100 kb, shifting the window by one SNP at the end of each step. Then one SNP from a pair of SNPs was removed if their LD was greater than 0.2. Both PCA and LD pruning were conducted in SNPRelate package in R software (Zheng et al., 2012). To investigate relatedness between individuals, a matrix of genomic relationship was calculated from marker data by the method described by (VanRaden, 2008) available in the R package snpReady (Granato et al., 2018).

### Statistical analysis of phenotypic data

Following a two-step approach, we initially obtained best linear unbiased estimates (BLUEs) of each genotype from analysis of individual environments. Note that in this first step the genotype effect was treated as fixed in order to prevent shrinkage in estimated means. BLUEs from this first step became the phenotype input for step two for combined analysis using a mixed model to estimate variance components, broad-sense heritability, and subsequent GWAS (Smith et al., 2001). The full description of analytical methods of multi-environment phenotypic data can be found in Supplemental Methods S3.

### Multi-environment GWAS analysis

For GWAS, we first extended the general mixed model form of the multi-environment analysis by adding genotype principal components into the fixed part of the model. In addition we incorporated the genomic relationships into the variance-covariance matrix of random effects to reflect the genetic relatedness between individuals in the population (Σ_*G*_ ⨂ *K*), allowing a diagonal residual matrix (different residual variances in each trial). GWAS was performed using the method proposed by Korte et al., (2012) which can be extended to multi-environment trials to identify QTL/SNPs either with main or interaction effects. The full description of analytical methods of multi-environment GWAS can be found in Supplemental Methods S4.

### Analysis of co-association network between traits

For each panel, we first organized associations from all traits into a matrix with SNPs (SNPs within the same LD region were treated as a single QTL) in rows and traits in columns and filled with cells for corresponding marker effects and its association with corresponding trait (QM, QF and interaction effects) after correction for population structure and kinship. The resulting matrix were then used to provide a pairwise Pearson correlations matrix between loci. The correlation matrix was subsequently used as an input matrix for network analysis. We used undirected graph networks to visualize submodules of loci using igraph package in R to visualize proximities between loci in a network plot (Csardi and Nepusz, 2006). Nodes (SNPs) were connected by edges if they had a pairwise correlation above threshold (r>= 0.9) from the similarity matrix described above.

## Supplemental Data

**Figure S1.**
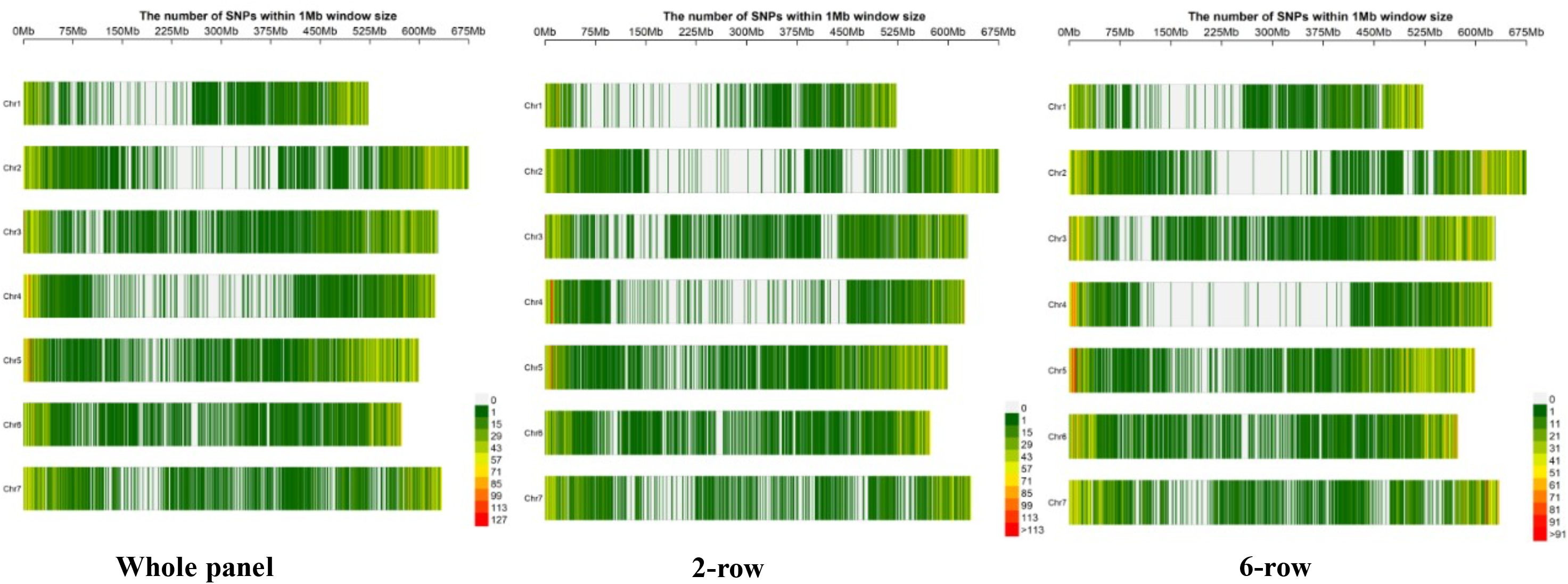
Distribution of SNP markers and marker density within the window size of 1Mb within the whole panel, 2-row, and 6-row panels, respectively. The number of markers per each chromosome are shown in the Supplemental Table S3.

**Figure S2.**
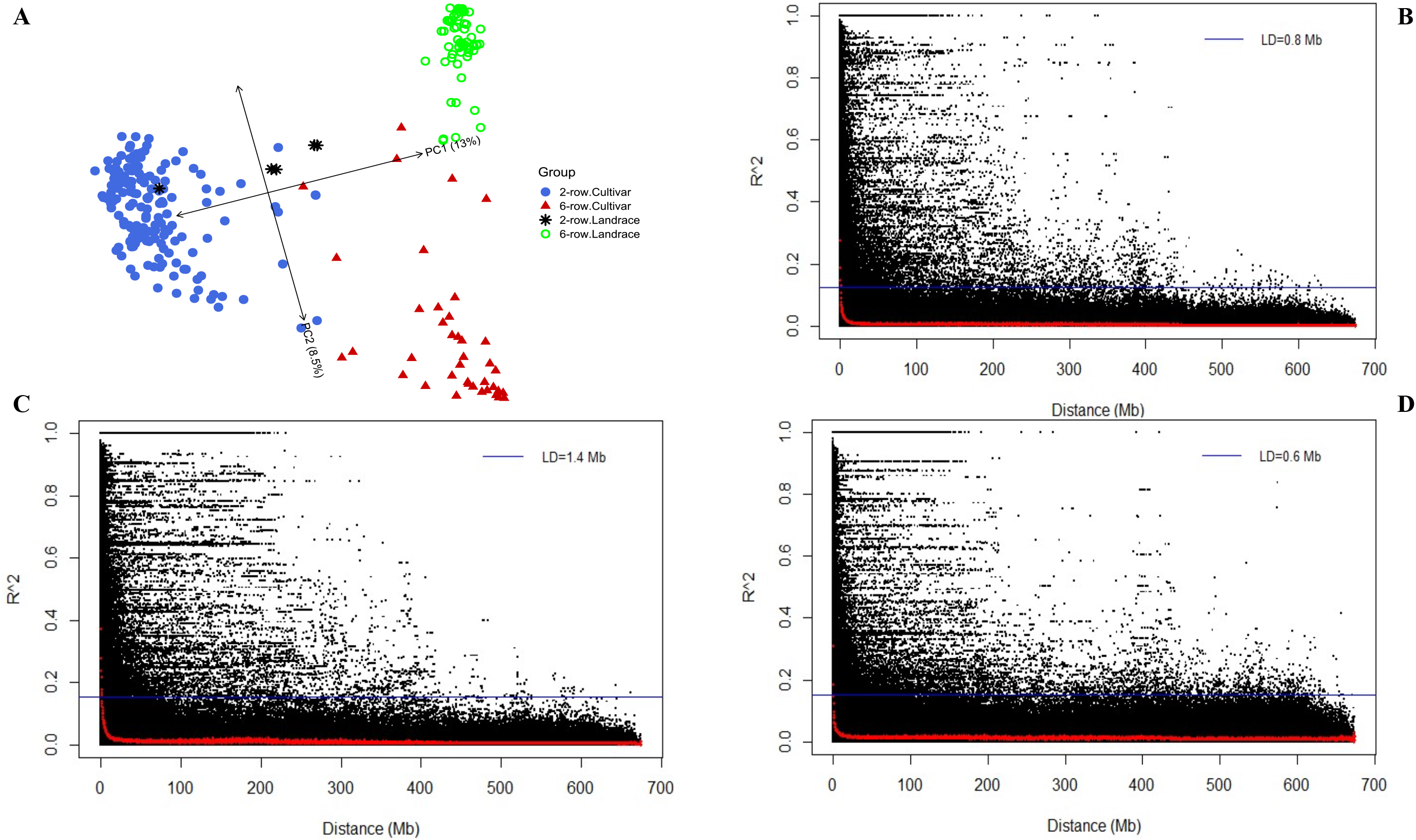
A) Biplot of first two PC scores from PCA analysis conducted on genotype marker data representing population structure of the panel related to row type and germplasm resource. Plots of LD (r^2^) decay corrected for population structure and relatedness representing intrachromosomal decay of marker pairs over all chromosomes as a function of physical distance. The blue line is the 95th percentile distribution of unlinked r^2^ values > 50 Mb and the red line illustrates the LD decay based on LOESS fitting curve. B) Whole panel; C) two-row panel; D) six-row panel.

**Figure S3.**
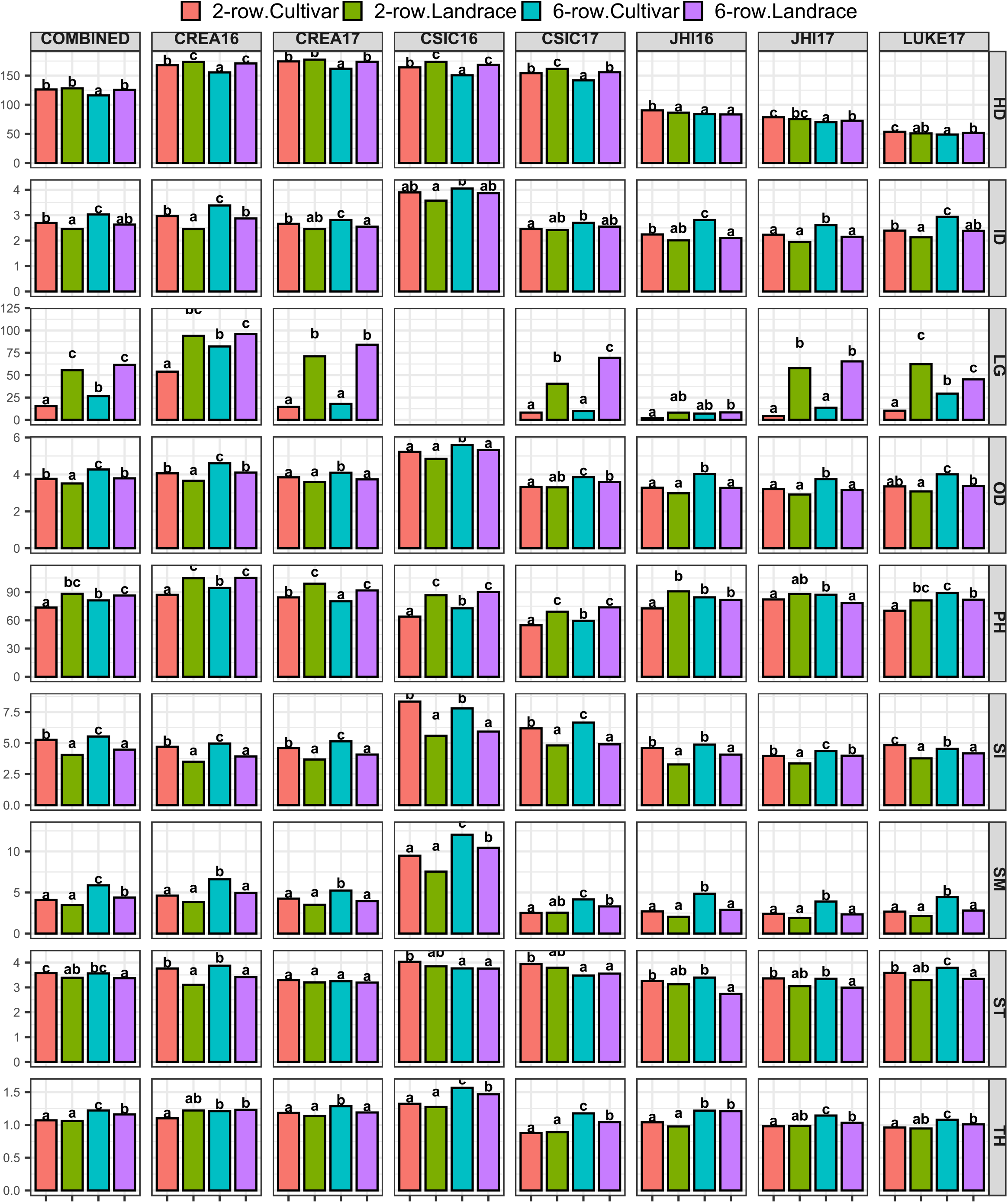
Comparison of row type and germplasm source for their effect on studied traits based on single- and across-environment trials within the barley panel. Mean differences were performed using a one-way ANOVA with Tukey’s honestly (HSD) test. Different letters above each column indicate significant differences (p-value= 0.05).

**Figure S4.**
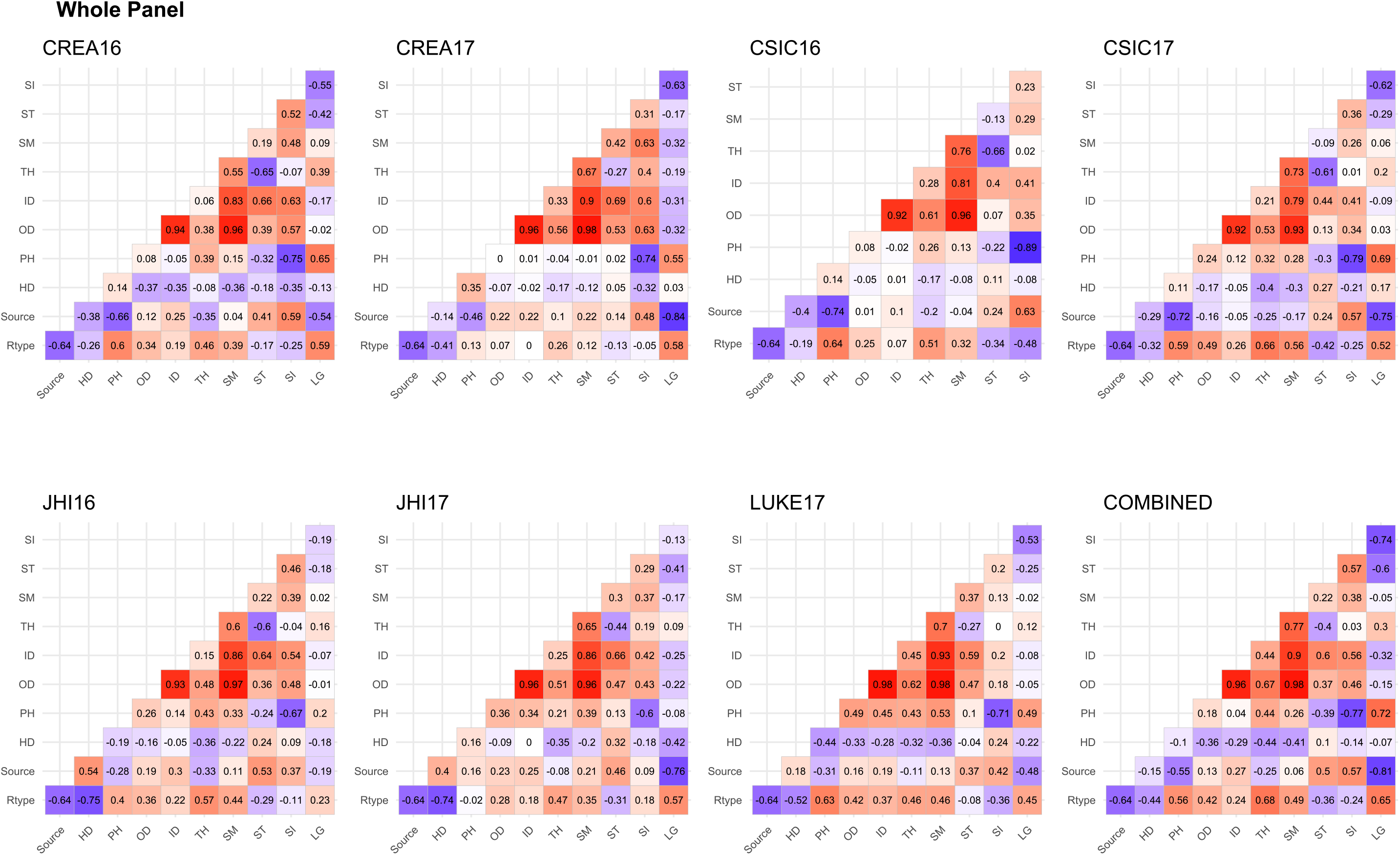
Pairwise correlation coefficients between traits, row type, and germplasm source (cultivar/landrace) in the whole panel based on genotype values estimated both in single and combined multi-environment analysis. Data for lodging In CSIC16 is not available.

**Figure S5.**
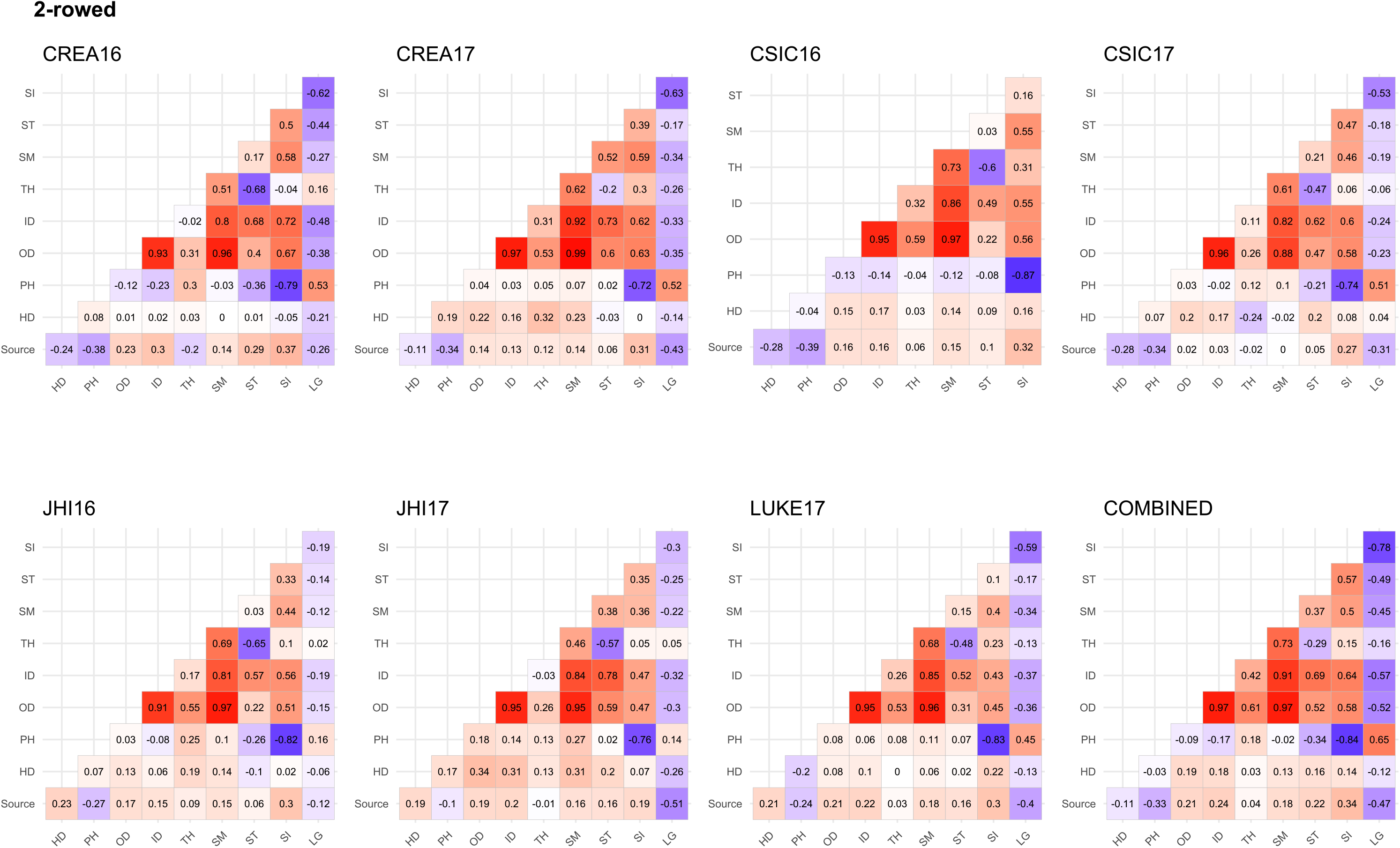
Pairwise correlation coefficients between traits and germplasm source (cultivar/landrace) in the 2-row panel based on genotype values estimated both in single and combined multi-environment analysis. Data for lodging In CSIC16 is not available.

**Figure S6.**
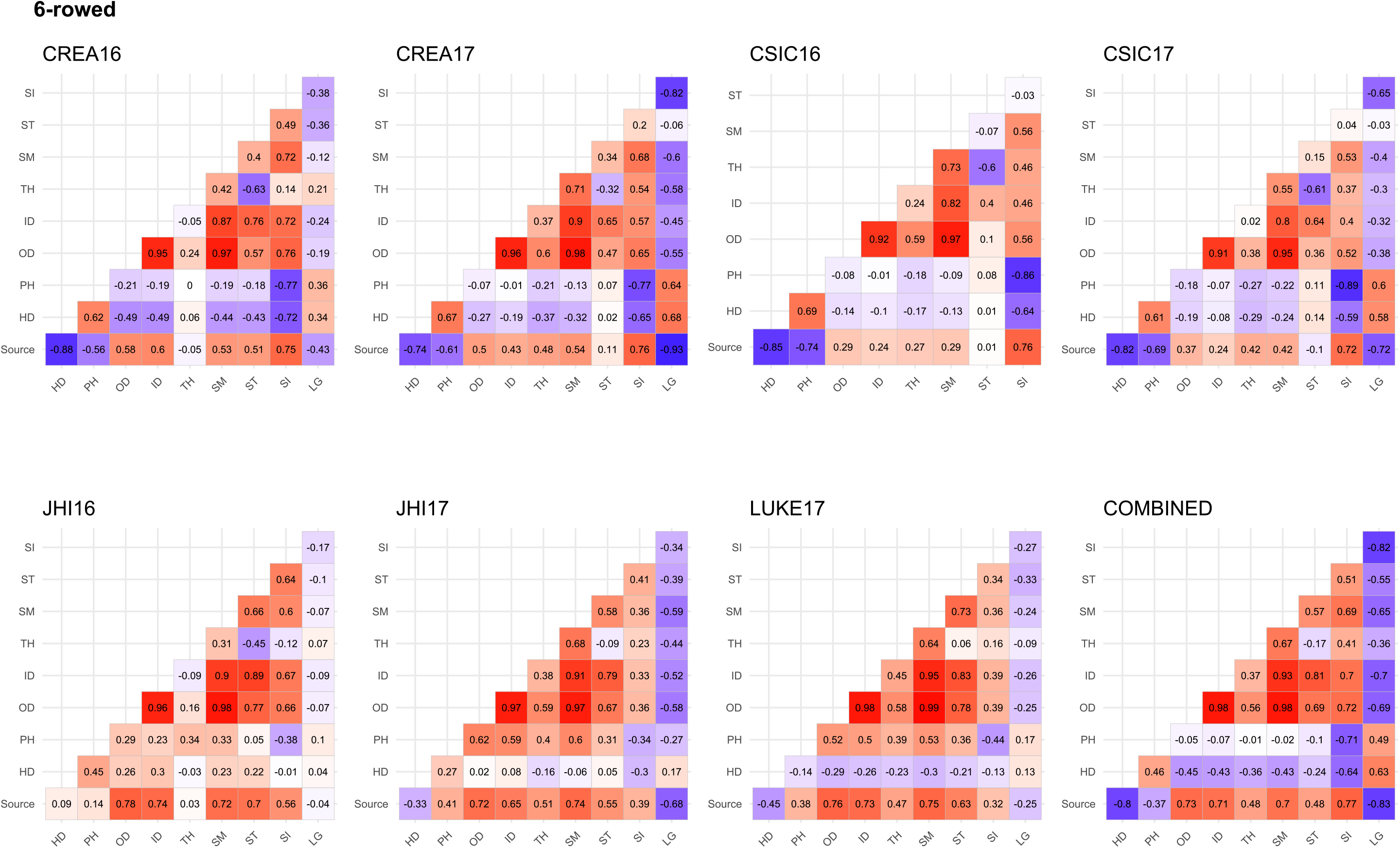
Pairwise correlation coefficients between traits and germplasm source (cultivar/landrace) in the 6-row panel based on genotype values estimated both in single and combined multi-environment analysis. Data for lodging In CSIC16 is not available.

**Figure S7.**
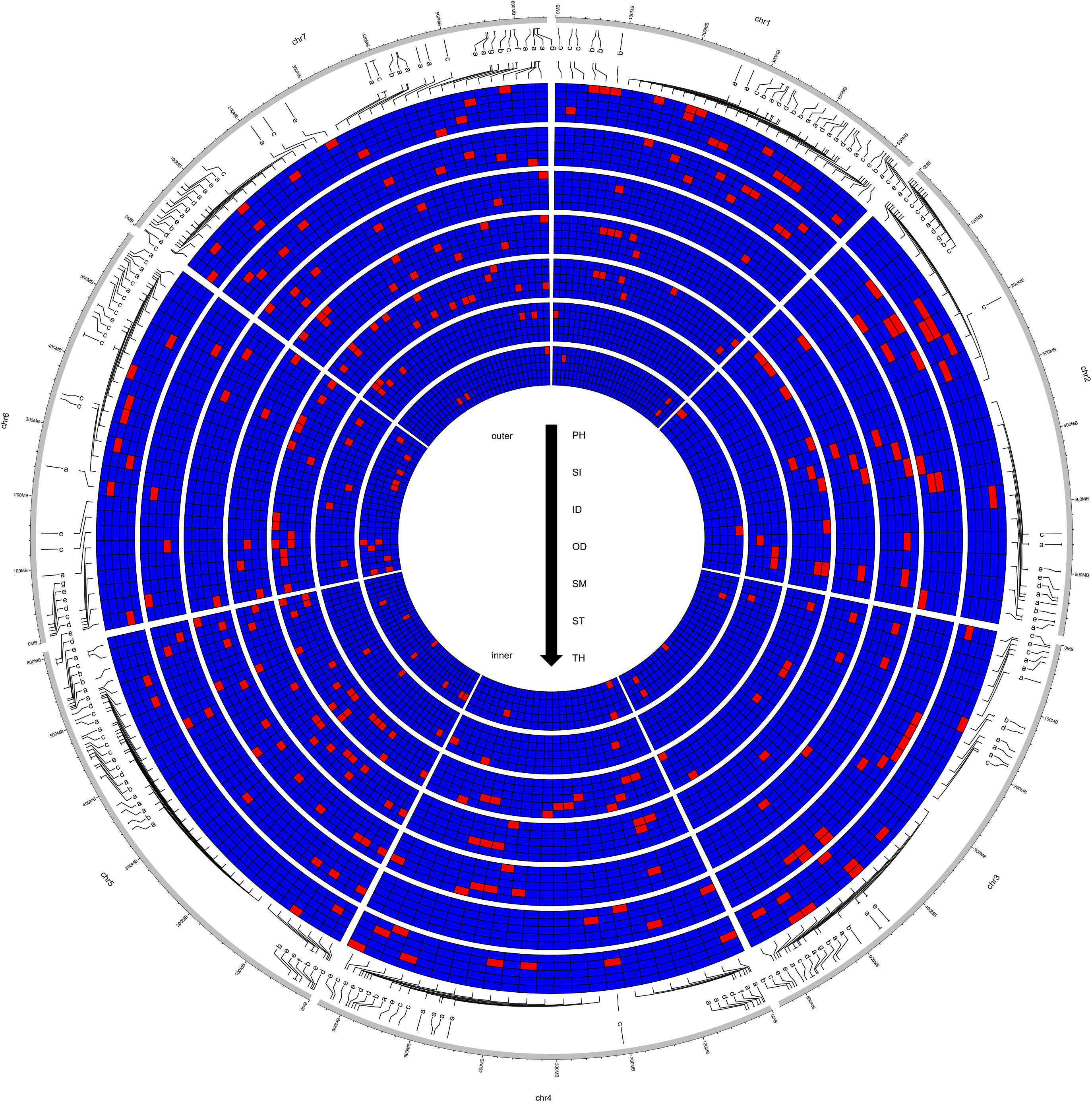
Circos heatmap for the 192 QTLs identified from GWAS of seven traits for the whole panel and row type groups. Each track belongs to one trait which also divided into five subsectors for QF, QM, QE, QL, and QY effect with red colors showing the presence of QTL at that position. The letters a, b, c, d, e, and f, are respectively related to QTLs identified in whole panel (a), 2-row (b), 6-row (d), both whole panel and 2-row (e), both whole panel and 6-row (f), and both 2-row and 6-row (g), and all the panels (h).

**Figure S8.**
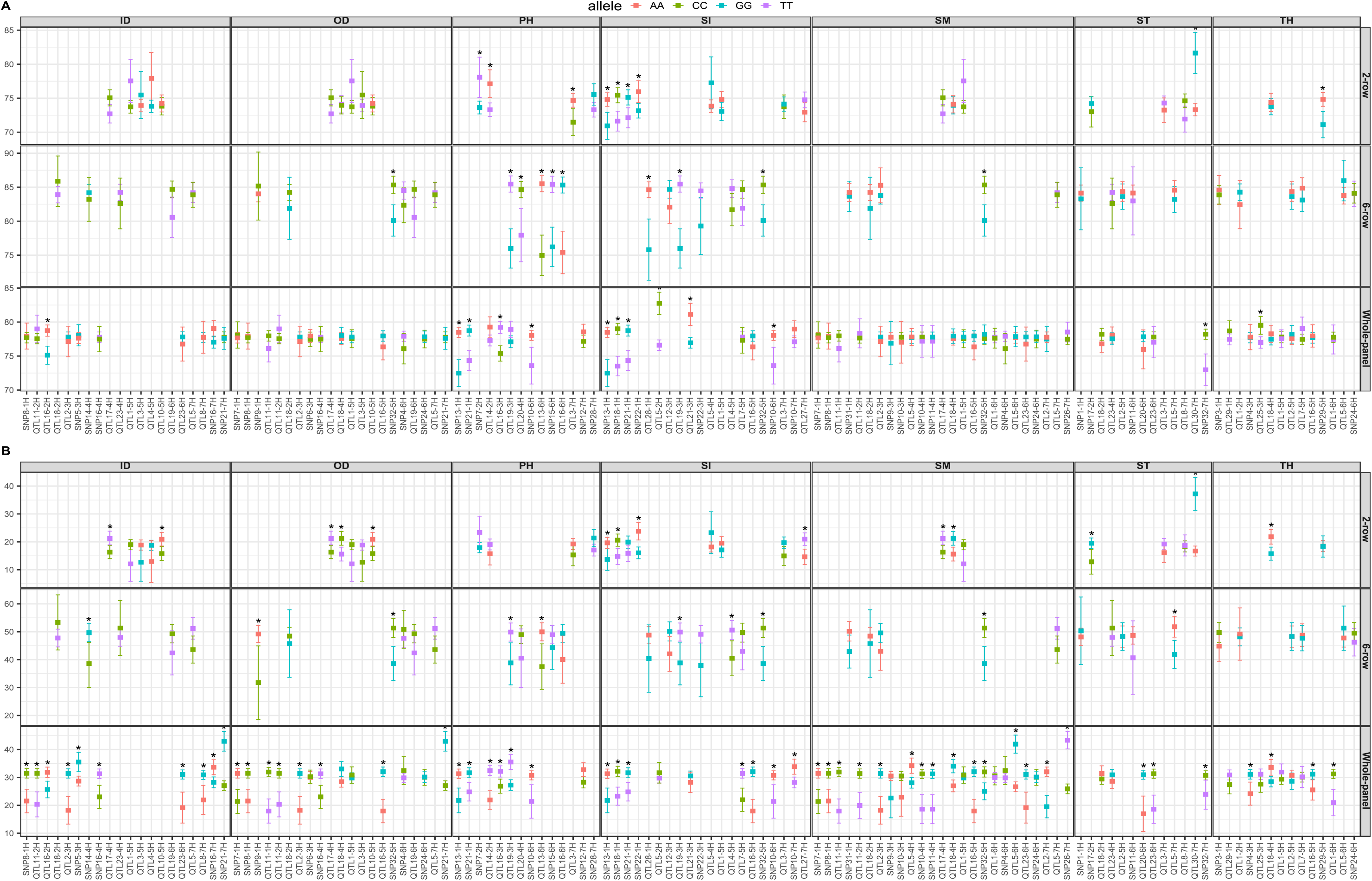
Comparison of allelic variants at peak markers of loci with QM effect (Supplemental Table S5). **A**) comparison between alleles at each marker for their effect on plant height; **B**) and their effect on lodging. The points indicate the mean value and the bars indicate the 95% confidence interval of the mean of corresponding allele. Significant differences are shown with asterisk.

**Figure S9.**
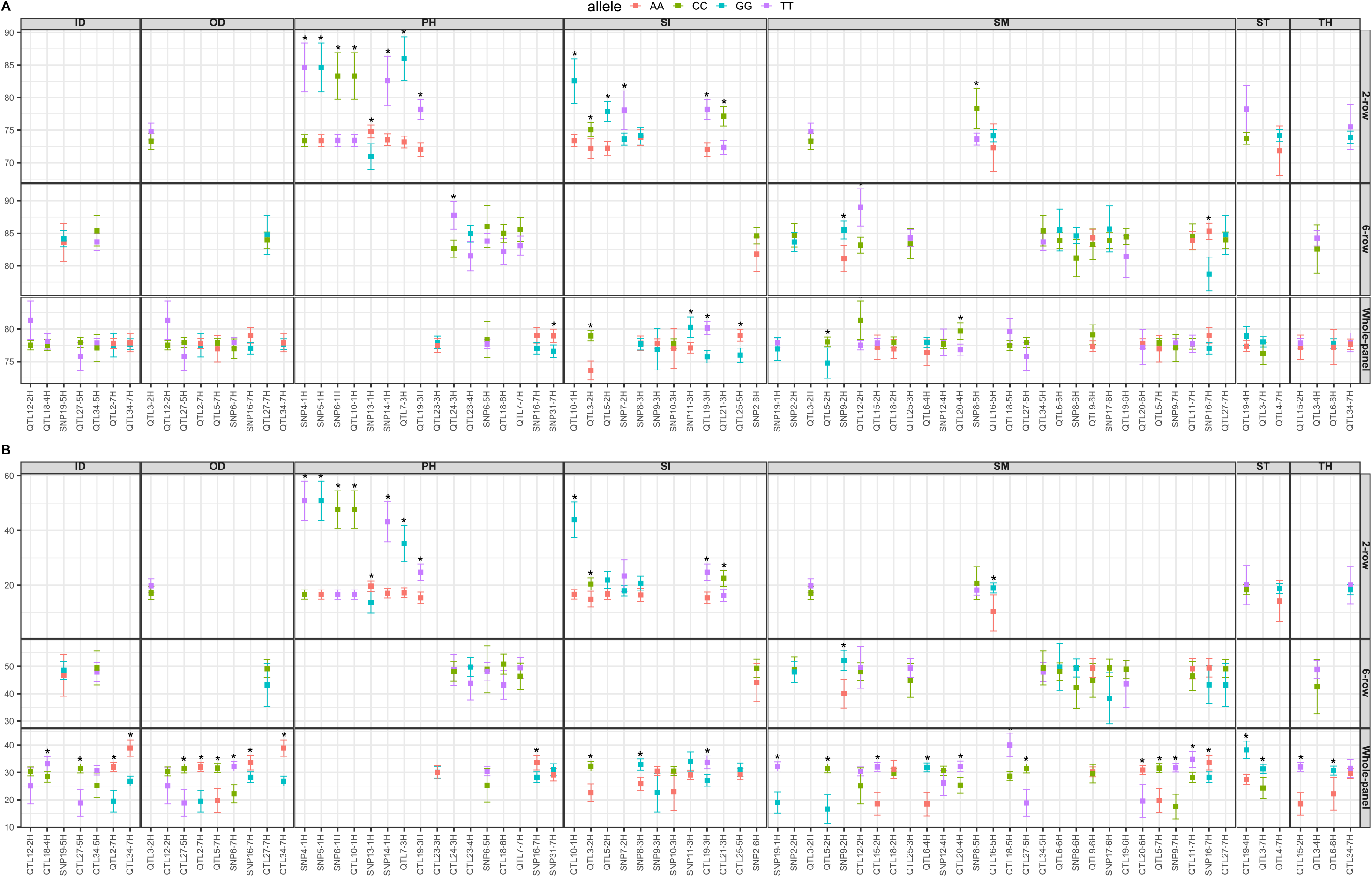
Comparison of allelic variants at peak markers of loci with QF effect (Supplemental Table S5). **A**) comparison between alleles at each marker for their effect on plant height; **B**) and their effect on lodging. The points indicate the mean value and the bars indicate the 95% confidence interval of the mean of corresponding allele. Significant differences are shown with asterisk.

**Supplemental Table S1**. List of 261 barley genotypes and their phenotypic information.

**Supplemental Table S2**. Number of SNP markers by chromosome.

**Supplemental Table S3**. Summary of experimental locations with phenotypic data.

**Supplemental Table S4**. Description of the traits and phenotyping methods measured in the panel.

**Supplemental Table S5**. Summary statistics of the eight traits measured in the 261 barley lines.

**Supplemental Table S6**. Complete list of QTLs detected in multi-environment GWAS.

## Acknowledgements

We are grateful to several students from University of Milan who helped in collecting phenotypic data. SS was in part supported by a post-doctoral fellowship (assegno di tipo A) from University of Milan.

## Methods S1

Additional methodological details on improved protocol for barley culm morphological traits.

Samples were collected at Zadoks stage 90 from the second internode of barley main culm, which is considered to different extent a critical area for lodging resistance (Pinthus 1974, Berry et al., 2004). Furthermore samples at Zadoks stage 90 have a different and more uniform cellular composition compared with other growth stages (Luo et al., 2007; Wang et al., 2018). The second basal internode was identified as the internode following the first internode longer than 1 cm above the root crown (Berry et al., 2004; Berry et al., 2007; Berry, 2013).

At Zadock stage 90 (fully mature), three randomly selected plants for each plot were uprooted, avoiding those on the plot’s borders. For each plant, the main stem was identified and the second basal internode was excised. Using a custom-made circular saw, internodes were cut in the central position to produce 5 mm thick cross-sections, taking care to produce blunt cuts. The resulting internode sections were attached with cyanoacrylic glue (Super Attak) to a black A4 cardboard, previously divided into 3cm x 5cm cells, each corresponding to a filed plot. Three samples (each from a distinct plant from the same plot) were glued in the same cell. On the side of each cardboard a paper ruler was attached in order to allow the software calibration during image analysis. Each section was then colored with a white marker (Uni-ball Posca, 0,7 mm) to ensure maximum contrast with the black background. In order to extract accurate measurements from culm sections, we developed a high-throughput image analysis protocol based on images obtained by scanning cardboards with a flat office scanner (600 dpi images in .tiff format).

The images were then analyzed to derive culm diameter and thickness data with a custom made macro command in Java language on the software ImageJ (Schindelin et al., 2012).

## Methods S2

Additional methodological details on missing genotype imputation.

To increase detection power and minimizing the loss of significant association, missing data were imputed using Beagle v5.0, which enables haplotypes inference and imputation of missing genotypes (Browning et al., 2018). Beagle uses a hidden Markov model to find the most likely haplotype pair for each individual given the genotype data for that individual. To estimate genotype phase the program works iteratively using an expectation –maximization method. Out of these markers, markers in perfect Linkage disequilibrium (LD) with adjacent SNP within the window size of 100kb (LD=1) were removed. Thus, a total of 33342 (Whole panel), 26262 (two-row subset), and 27583 (six-row subset) SNPs were left for calculation of kinship matrix and subsequent GWAS analysis.

## Methods S3

Additional methodological details on statistical analysis of phenotypic data, computation of adjusted means, estimation of variance components, and heritability.

Following a two-stage approach, in each environment (stage 1) with two replicates and coordinates of column and rows, a mixed model was used. We treated the genotype (to obtain BLUEs) and replicate as fixed effects and row and columns as random effects. Depending of the trial, the residual effects were also modelled using spatial methods that accommodate local or plot to plot variation (Table 1).

After calculation of BLUEs for each trait from all seven environments, approximately 6% of data mainly from six-row panel were missing in the genotype-environment table (i.e, for three environments JHI16, JHI17, and LUKE17). Removing accessions with missing phenotype will reduce the sample size and consequently will negatively impact the statistical inference and subsequent GWAS analysis (Rodrigues et al., 2014; Scutari et al., 2014; Dahl et al., 2016).

Therefore, prior to subsequent analysis, and due to small fraction of missing phenotypes existed in our data, we performed imputation of missing cells in a genotype-by-environment table using the Expectation Maximization Additive Main Effects and Multiplicative Interaction (EM-AMMI) algorithm (Cauch and Zobel, 1990; Gauch 1992). We run the algorithm using five steps with the R script indicated as follows (Cauch and Zobel, 1990; Paderewski and Rodrigues, 2014): At first, initial values were assigned to missing cells; secondly, the parameters of the AMMI model were estimated; third, the adjusted means were calculated according to principal components obtained from AMMI analysis; next, missing cells were filled based on adjusted means and ;finally, the steps from 2 to 5 were repeated if the Chebyshev distance between the missing value estimations in the two progressive iteration steps were greater than the assumed precision, otherwise the algorithm was stopped. We considered the results as reliable, as the relationships between the genotypes and environments for almost all traits were present. The important factor of the algorithm is to select appropriate number of principal components to be included in imputation process. We selected this number based on the minimum of the Root Mean Square Predictive Difference (RMSPD, Gauch and Zobel, 1990; Dias and Krzanowski, 2003). The appropriate number of principal components is the one with the smallest RMSPD value. The RMSPD values were calculated according to leave-one-out cross validation (LOO-CV) procedure. Briefly, a single non-missing phenotype is hidden from the dataset and EM-AMMI is employed on training data (without missing). The procedure is repeated for each observation until no empty cell remained in the dataset. The RMSPD, is then obtained based on the difference between the hidden value and the value imputed by EM-AMMI (the predictive differences). We initially performed association analysis both on imputed data and the data after removing missing cells and found that, although the results were highly similar, the analysis with imputed phenotypes, in accordance with previous studies, resulted in well-calibrated p-values due to increased sample size (Scutari et al., 2014; Dahl et al., 2016).

In stage 2, the resulting BLUEs were used for combined analysis using a mixed model to estimate variance components, broad-sense heritability, and subsequent GWAS. Variance components and heritability values were estimated under the general form of mixed model:

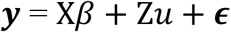

where y is a vector of observations (phenotypic BLUEs across environments), X is the design matrix for fixed effects β (intercepts and environment), Z is the design matrix for random effects (genotypes). *u* is the vector of random effects with *u*∼*N*(*o*. **Σ**_***G***_) and ϵ∼*N*(*o*. ***R***). The **Σ**_***G***_ is between environment variance-covariance matrix and **R** is a diagonal block matrix where :

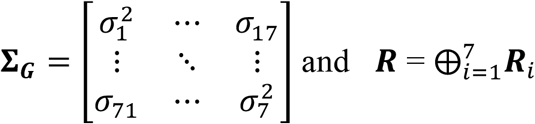

The specification of variance structure is important in combined analysis. Traditionally the genotypic variances within all environments and the covariances between genotypic values for each pair of environments are assumes equal. We relaxed these assumptions for **Σ**_***G***_ using the mixed model allowing for unequal genotype variances and unique covariances for each pair of environments. Therefore we specified the unstructured covariance and heterogeneous variance (US) model in the multi-environment analysis (7 within-environment variances and 14 between-environment covariances). The genotype means from combined multi-environment analysis (BLUPs) were then obtained for comparisons with single environments and for correlation analysis between traits.

Using average covariance between genotypes across environments as the numerator and average variance of genotype means across environments as the denominator we estimated the heritability using the following formula:

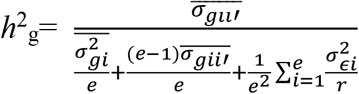

where ***e*** refers to the number of environments, ***r*** refers to the number of replications within environment, *σg*_*ii*_, is the genotype covariance between environments i and i’, 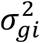 is the genotype variance within environment i, and 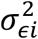 is assumed to follow ϵ_*i*_∼*N*(*o*. ***R***). in environment i and 0 is diagonal matrix calculated from squared errors of genotype BLUEs from stage 1. If the covariance between environments is higher, the heritability would be high accordingly. The variance parameters were estimated by maximizing the REML (Patterson and Thompson, 1971) log-likelihood function using the AI algorithm (Gilmour et al., 1995), implemented in the package ASReml-R (Butler et al., 2017). Pairwise correlations between traits based on genotype means estimated from each environment and across environments were calculated using R package ggcorrplot.

## Methods S4

Additional methodological details on multi-environment GWAS analysis.

The MTMM can be written as follow:

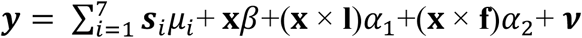

Where ***y*** is the vector of phenotypic BLUEs across environments, ***x*** is the vector of marker scores and ***s***_*i*_ is a vector having 1 for values belonging to the i’th environment and 0 otherwise. **l** is a vector with 1 for all the values measured in the same location, **f** is a vector with 1 for all the values measured in the same year, and ***v*** ∼ *N*(*o*. **Σ**_***G***_ ⨂ ***K*** + ***R***) is a random variable comprising of both residual and random genetic effects. The variance of ***v*** is estimated from a variance decomposition model described above. A generalized least square (GLS) F-test was used to estimate the genome-wide marker effects depending on what kind of QTL/SNP we were interested as follows:

**QF**, This is the full model which tested against null model *β*= *α*_1_=*α*_2_=0 which identifies SNPs with both stable and interaction effects;

**QM**, To identify the main QTL which tests the model with *α*_1_=*α*_2_=0 against the null model with *β*=*α*_1_=*α*_2_=0;

**QL**, To identify the QTL × location interaction which tests the full model against the null model with *α*_1_=0.

**QY**: To identify the QTL × year interaction the full model tested against null model with *α*_2_=0.

**QE**: To identify any QTL × environment interaction effect where the full model is tested against null model with *α*_1_= *α*_2_=0.

For marker-trait association, we didn’t use the Bonferroni adjustment due to its highly conservative nature and overcorrect for SNPs falling in high linkage disequilibrium that are not truly independent. Therefore, we approximated GWAS p-value significance thresholds according to the true number of ‘independent SNP tests’. This effective number of SNPs was estimated in software Haploview 4.2 (Barret et al., 2005) using r-square tag threshold estimated from LD decay analysis (see LD section) (Mackay, 1996). We also retained the associations with –log_10_ P ≥ 4 but lower than the significance threshold as suggestive QTLs. Haploview was also used to determine the extent of QTL intervals within the barley chromosomes where SNPs detected in the same haplotype blocks were considered as the same QTL. To estimate the proportion of phenotypic variance explained by an SNP, we were faced with either a single SNP or multiple SNPs in the region with high LD between them. In the case of first situation we calculated variance using the following formulae: *CDEα*(%) = 2*p*_i_(1-*p*_i_)β ×100 where β is the main effect derived from the GWAS model and *p*_i_ is the frequency of minor allele at SNP_i_. In the case of QTL region with multiple associated SNPs, the phenotypic variance explained by the QTL was calculated as: 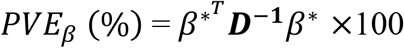, where *β*^∗^ is a matrix with the elements 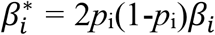 and 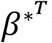 is the transposed matrix. **D** Is the LD-matrix (Pearson correlations) of the variants in the QTL region. To derive PVE (%) explained by QTL-by-Location and QTL-by-Year effects, the *β*_*i*_ was replaced by *α*_1_ and *α*_2_, respectively. Finally, the total phenotype variance was obtained by summing over main and interaction effects.

## Literature cited

Agnew E, Bray A, Floro E, Ellis N, Gierer J, Lizárraga C, O’Brien D, Wiechert M, Mockler TC, Shakoor N, et al (2017) Whole-plant manual and image-based phenotyping in controlled environments. Curr Protoc Plant Biol 2: 1–21

Aya K, Hobo T, Sato-Izawa K, Ueguchi-Tanaka M, Kitano H, Matsuoka M (2014) A novel AP2-type transcription factor, SMALL ORGAN SIZE1, controls organ size downstream of an auxin signaling pathway. Plant Cell Physiol 55: 897–912

Bayer MM, Rapazote-Flores P, Ganal M, Hedley PE, Macaulay M, Plieske J, Ramsay L, Russell J, Shaw PD, Thomas W, et al (2017) Development and evaluation of a barley 50k iSelect SNP array. Front Plant Sci 8: 1792

Berry PM (2013) Lodging resistance in cereals. In P Christou, R Savin, BA Costa-Pierce, I Misztal, CBA Whitelaw, eds, Sustain. Food Prod. Springer New York, New York, NY, pp 1096–1110

Berry PM, Spink J (2012) Predicting yield losses caused by lodging in wheat. Field Crop Res 137: 19–26

Berry PM, Sterling M, Spink JH, Baker CJ, Sylvester-Bradley R, Mooney SJ, Tams AR, Ennos AR (2004) Understanding and reducing lodging in cereals. Adv Agron 84: 217–271

Breseghello F, Sorrells ME (2006) Association mapping of kernel size and milling quality in wheat (Triticum aestivum L.) cultivars. Genetics 172: 1165–1177

Browning BL, Zhou Y, Browning SR (2018) A one-penny imputed genome from next-generation reference panels. Am J Hum Genet 103: 338–348

Burton RA, Ma G, Baumann U, Harvey AJ, Shirley NJ, Taylor J, Pettolino F, Bacic A, Beatty M, Simmons CR, et al (2010) A customized gene expression microarray reveals that the brittle stem phenotype fs2 of barley Is attributable to a retroelement in the HvCesA4 cellulose synthase gene. Plant Physiol 153: 1716–1728

Cai T, Peng D, Wang R, Jia X, Qiao D, Liu T, Jia Z, Wang Z, Ren X (2019) Can intercropping or mixed cropping of two genotypes enhance wheat lodging resistance? Field Crop Res 239: 10–18

Chandler PM, Harding CA (2013) ‘Overgrowth’ mutants in barley and wheat: new alleles and phenotypes of the ‘Green Revolution’ DELLA gene. J Exp Bot 64: 1603–1613

Csardi G, Nepusz T (2006) The igraph software package for complex network research. Inter Journal Complex Syst 1695:1–9

Chesterfield RJ, Vickers CE, Beveridge CA (2020) Translation of strigolactones from plant hormone to agriculture: achievements, future perspectives, and challenges. Trends Plant Sci 25: 1087–1106

Chono M, Honda I, Zeniya H, Yoneyama K, Saisho D, Takeda K, Takatsuto S, Hoshino T, Watanabe Y (2003) A semidwarf phenotype of barley uzu results from a nucleotide substitution in the gene encoding a putative brassinosteroid receptor. Plant Physiol 133: 1209–1219

Cui Y, Hu X, Liang G, Feng A, Wang F, Ruan S, Dong G, Shen L, Zhang B, Chen D, et al (2020) Production of novel beneficial alleles of a rice yield-related QTL by CRISPR/Cas9. Plant Biotechnol J 18: 1987–1989

Dahleen LS, Vander Wal LJ, Franckowiak JD (2005) Characterization and molecular mapping of genes determining semidwarfism in barley. J Hered 96: 654–662

Dawson IK, Russell J, Powell W, Steffenson B, Thomas WTB, Waugh R (2015) Barley: a translational model for adaptation to climate change. New Phytol 206: 913–931

Digel B, Tavakol E, Verderio G, Tondelli A, Xu X, Cattivelli L, Rossini L, von Korff M (2016) Photoperiod-H1 (Ppd-H1) controls leaf size. Plant Physiol 172: 405–415

Dockter C, Gruszka D, Braumann I, Druka A, Druka I, Franckowiak J, Gough SP, Janeczko A, Kurowska M, Lundqvist J, et al (2014) Induced variations in brassinosteroid genes define barley height and sturdiness, and expand the green revolution genetic toolkit. Plant Physiol 166: 1912–1927

Duan K, Li L, Hu P, Xu SP, Xu ZH, Xue HW (2006) A brassinolide-suppressed rice MADS-box transcription factor, OsMDP1, has a negative regulatory role in BR signaling. Plant J 47: 519–531

Göransson M, Hallsson JH, Lillemo M, Orabi J, Backes G, Jahoor A, Hermannsson J, Christerson T, Tuvesson S, Gertsson B, et al (2019) Identification of ideal allele combinations for the adaptation of spring barley to northern latitudes. Front Plant Sci 10: 542

Granato ISC, Galli G, de Oliveira Couto EG, e Souza MB, Mendonça LF, Fritsche-Neto R (2018) snpReady: a tool to assist breeders in genomic analysis. Mol Breed 38: 102

He T, Hill CB, Angessa TT, Zhang X-Q, Chen K, Moody D, Telfer P, Westcott S, Li C (2019) Gene-set association and epistatic analyses reveal complex gene interaction networks affecting flowering time in a worldwide barley collection. J Exp Bot 70: 5603–5616

Hemming MN, Peacock WJ, Dennis ES, Trevaskis B (2008) Low-temperature and daylength cues are integrated to regulate Flowering Locus T in barley. Plant Physiol 147: 355–366

Hernandez J, Meints B, Hayes P (2020) Introgression breeding in barley: perspectives and case studies. Front Plant Sci 11: 761

Hirano K, Okuno A, Hobo T, Ordonio R, Shinozaki Y, Asano K, Kitano H, Matsuoka M (2014) Utilization of stiff culm trait of rice smos1 mutant for increased lodging resistance. PLoS One 9: e96009

Hirano K, Ordonio RL, Matsuoka M (2017a) Engineering the lodging resistance mechanism of post-green revolution rice to meet future demands. Proc Japan Acad Ser B Phys Biol Sci 93: 220–233

Hirano K, Yoshida H, Aya K, Kawamura M, Hayashi M, Hobo T, Sato-Izawa K, Kitano H, Ueguchi-Tanaka M, Matsuoka M (2017b) SMALL ORGAN SIZE 1 and SMALL ORGAN SIZE 2/DWARF AND LOW-TILLERING form a complex to integrate auxin and brassinosteroid signaling in rice. Mol Plant 10: 590–604

Houston K, Burton RA, Sznajder B, Rafalski AJ, Dhugga KS, Mather DE, Taylor J, Steffenson BJ, Waugh R, Fincher GB (2015) A genome-wide association study for culm cellulose content in barley reveals candidate genes co-expressed with members of the CELLULOSE SYNTHASE A gene family. PLoS One 10: e0130890

Ilg A, Beyer P, Al-Babili S (2009) Characterization of the rice carotenoid cleavage dioxygenase 1 reveals a novel route for geranial biosynthesis. FEBS J 276: 736–747

Islam MS, Peng S, Visperas RM, Ereful N, Sultan M, Bhuiya U, Julfiquar AW (2007) Lodging-related morphological traits of hybrid rice in a tropical irrigated ecosystem. Field Crop Res 101: 240–248

Jedel PE, Helm JH (1991) Lodging effects on a semidwarf and two standard barley cultivars. Agron J 83: 158–161

Karsai I, Mészáros K, Szücs P, Hayes PM, Láng L, Bedö Z (1999) Effects of loci determining photoperiod sensitivity (Ppd-H1) and vernalization response (Sh2) on agronomic traits in the “Dicktoo” x “Morex” barley mapping population. Plant Breed 118: 399–403

Khush GS (2001) Green revolution: The way forward. Nat Rev Genet 2: 815–822

Komatsuda T, Pourkheirandish M, He C, Azhaguvel P, Kanamori K, Perovic D, Stein N, Graner A, Wicker T, Tagiri A, et al (2007) Six-rowed barley originated from a mutation in a homeodomain-leucine zipper I-class homeobox gene. Proc Natl Acad Sci U S A 104: 1424–1429

Korte A, Vilhjálmsson BJ, Segura V, Platt A, Long Q, Nordborg M (2012) A mixed-model approach for genome-wide association studies of correlated traits in structured populations. Nat Genet 44: 1066–1071

Kuczyńska A, Surma M, Adamski T, Mikołajczak K, Krystkowiak K, Ogrodowicz P (2013) Effects of the semi-dwarfing sdw1/denso gene in barley. J Appl Genet 54: 381–390

Kuczynska A, Wyka T (2011) The effect of the denso dwarfing gene on morpho-anatomical characters in barley recombinant inbred lines. Breed Sci 61: 275–280

Liller CB, Neuhaus R, von Korff M, Koornneef M, van Esse W (2015) Mutations in barley row type genes have pleiotropic effects on shoot branching. PLoS One 10: e0140246

Lin H, Wang R, Qian Q, Yan M, Meng X, Fu Z, Yan C, Jiang B, Su Z, Li J, et al (2009) DWARF27,an iron-containing protein required for the biosynthesis of strigolactones,regulates rice tiller bud outgrowth. Plant Cell 21: 1512–1525

Liu L, Tong H, Xiao Y, Che R, Xu F, Hu B, Liang C, Chu J, Li J, Chu C (2015) Activation of Big Grain1 significantly improves grain size by regulating auxin transport in rice. Proc Natl Acad Sci U S A 112: 11102–11107

Lobell DB, Schlenker W, Costa-Roberts J (2011) Climate trends and global crop production since 1980. Science (80-) 333: 616–620

Malik N, Ranjan R, Parida SK, Agarwal P, Tyagi AK (2020) Mediator subunit OsMED14_1 plays an important role in rice development. Plant J 101: 1411–1429

Mangin B, Siberchicot A, Nicolas S, Doligez A, This P, Cierco-Ayrolles C (2012) Novel measures of linkage disequilibrium that correct the bias due to population structure and relatedness. Heredity 108: 285–291

Mantilla Perez MB, Zhao J, Yin Y, Hu J, Salas Fernandez MG (2014) Association mapping of brassinosteroid candidate genes and plant architecture in a diverse panel of Sorghum bicolor. Theor Appl Genet 127: 2645–2662

Minakuchi K, Kameoka H, Yasuno N, Umehara M, Luo L, Kobayashi K, Hanada A, Ueno K, Asami T, Yamaguchi S, et al (2010) FINE CULM1 (FC1) works downstream of strigolactones to inhibit the outgrowth of axillary buds in rice. Plant Cell Physiol 51: 1127–1135

Monat C, Padmarasu S, Lux T, Wicker T, Gundlach H, Himmelbach A, Ens J, Li C, Muehlbauer GJ, Schulman AH, et al (2019) TRITEX: chromosome-scale sequence assembly of Triticeae genomes with open-source tools. Genome Biol 20: 284

Mulsanti IW, Yamamoto T, Ueda T, et al (2018) Finding the superior allele of japonica-type for increasing stem lodging resistance in indica rice varieties using chromosome segment substitution lines. Rice 11: 25

Okuno A, Hirano K, Asano K, Takase W, Masuda R, Morinaka Y, Ueguchi-Tanaka M, Kitano H, Matsuoka M (2014) New approach to increasing rice lodging resistance and biomass yield through the use of high gibberellin producing varieties. PLoS One 9: e86870

Ookawa T, Hobo T, Yano M, Murata K, Ando T, Miura H, Asano K, Ochiai Y, Ikeda M, Nishitani R, et al (2010) New approach for rice improvement using a pleiotropic QTL gene for lodging resistance and yield. Nat Commun 1: 132

Peng J, Richards DE, Hartley NM, Murphy GP, Devos KM, Flintham JE, Beales J, Fish LJ, Worland AJ, Pelica F, et al (1999) “Green revolution” genes encode mutant gibberellin response modulators. Nature 400: 256–261

Piñera-Chavez FJ, Berry PM, Foulkes MJ, Molero G, Reynolds MP (2016) Avoiding lodging in irrigated spring wheat. II. genetic variation of stem and root structural properties. Field Crop Res 196: 64–74

Pinthus MJ (1974) Lodging in wheat, barley, and oats: the phenomenon, its causes, and preventive measures. Adv Agron 25: 209–263

Priya R, Siva R (2014) Phylogenetic analysis and evolutionary studies of plant carotenoid cleavage dioxygenase gene. Gene 548: 223–233

Qiao S, Sun S, Wang L, Wu Z, Li C, Li X, Wang T, Leng L, Tian W, Lu T, et al (2017) The RLA1/SMOS1 transcription factor functions with OsBZR1 to regulate brassinosteroid signaling and rice architecture. Plant Cell 29: 292–309

Qiao Y, Piao R, Shi J, Lee SI, Jiang W, Kim BK, Lee J, Han L, Ma W, Koh HJ (2011) Fine mapping and candidate gene analysis of dense and erect panicle 3, DEP3, which confers high grain yield in rice (Oryza sativa L.). Theor Appl Genet 122: 1439–1449

Rajkumara S (2008) Lodging in cereals-a review. Agric Rev 29: 55–60

Ramsay L, Comadran J, Druka A, Marshall DF, Thomas WTB, MacAulay M, MacKenzie K, Simpson C, Fuller J, Bonar N, et al (2011) INTERMEDIUM-C, a modifier of lateral spikelet fertility in barley, is an ortholog of the maize domestication gene TEOSINTE BRANCHED 1. Nat Genet 43: 169–172

Rockström J, Williams J, Daily G, Noble A, Matthews N, Gordon L, Wetterstrand H, DeClerck F, Shah M, Steduto P, et al (2017) Sustainable intensification of agriculture for human prosperity and global sustainability. Ambio 46: 4–17

Samadi AF, Suzuki H, Ueda T, Yamamoto T, Adachi S, Ookawa T (2019) Identification of quantitative trait loci for breaking and bending types lodging resistance in rice, using recombinant inbred lines derived from Koshihikari and a strong culm variety, Leaf Star. Plant Growth Regul 89: 83–98

Sameri M, Nakamura AS, Nair ASK, Ae AKT, Komatsuda T (2009) A quantitative trait locus for reduced culm internode length in barley segregates as a Mendelian gene. Theor Appl Genet 118: 643–652

Sasaki A, Ashikari M, Ueguchi-Tanaka M, Itoh H, Nishimura A, Swapan D, Ishiyama K, Saito T, Kobayashi M, Khush GS, et al (2002) A mutant gibberellin-synthesis gene in rice: new insight into the rice variant that helped to avert famine over thirty years ago. Nature 416: 701–702

Shah L, Yahya M, Shah SMA, Nadeem M, Ali A, Ali A, Wang J, Riaz MW, Rehman S, Wu W, et al (2019) Improving lodging resistance: using wheat and rice as classical examples. Int J Mol Sci 20: 4211

Shi C, Ren Y, Liu L, Wang F, Zhang H, Tian P, Pan T, Wang Y, Jing R, Liu T, et al (2019) Ubiquitin specific protease 15 has an important role in regulating grain width and size in rice. Plant Physiol 180: 381–391

Smith A, Cullis B, Gilmour A (2001) The analysis of crop variety evaluation data in Australia. Aust New Zeal J Stat 43: 129–145

Sowadan O, Li D, Zhang Y, Zhu S, Hu X, Bhanbhro LB, Edzesi WM, Dang X, Hong D (2018) Mining of favorable alleles for lodging resistance traits in rice (Oryza sativa) through association mapping. Planta 248: 155–169

Sui P, Jin J, Ye S, Mu C, Gao J, Feng H, Shen W-H, Yu Y, Dong A (2012) H3K36 methylation is critical for brassinosteroid-regulated plant growth and development in rice. Plant J 70: 340–347

Tondelli A, Xu X, Moragues M, Sharma R, Schnaithmann F, Ingvardsen C, Manninen O, Comadran J, Russell J, Waugh R, et al (2013) Structural and temporal variation in genetic diversity of european spring two-row barley cultivars and association mapping of quantitative traits. Plant Genome. 6: 1–14

Tong H, Jin Y, Liu W, Li F, Fang J, Yin Y, Qian Q, Zhu L, Chu C (2009) DWARF AND LOW-TILLERING, a new member of the GRAS family, plays positive roles in brassinosteroid signaling in rice. Plant J 58: 803–816

Tong H, Liu L, Jin Y, Du L, Yin Y, Qian Q, Zhu L, Chu C (2012) DWARF AND LOW-TILLERING acts as a direct downstream target of a GSK3/SHAGGY-Like kinase to mediate brassinosteroid responses in rice. Plant Cell 24: 2562–2577

Turner A, Beales J, Faure S, Dunford RP, Laurie DA (2005) The pseudo-response regulator Ppd-H1 provides adaptation to photoperiod in barley. Science 310: 1031–1034

VanRaden PM (2008) Efficient methods to compute genomic predictions. J Dairy Sci 91: 4414–4423

Walla A, Wilma van Esse G, Kirschner GK, Guo G, Brünje A, Finkemeier I, et al (2020) An acyl-CoA N-acyltransferase regulates meristem phase change and plant architecture in barley. Plant Physiol 183: 1088–109

Xu Y, Jia Q, Zhou G, Zhang X-Q, Angessa T, Broughton S, Yan G, Zhang W, Li C (2017) Characterization of the sdw1 semi-dwarf gene in barley. BMC Plant Biol 17: 11

Yano K, Ookawa T, Aya K, Ochiai Y, Hirasawa T, Ebitani T, Takarada T, Yano M, Yamamoto T, Fukuoka S, et al (2015) Isolation of a novel lodging resistance QTL gene involved in strigolactone signaling and its pyramiding with a QTL gene involved in another mechanism. Mol Plant 8: 303–314

Zadoks JC, Chang TT, Konzak CF (1974) A decimal code for the growth stages of cereals. Weed Res 14: 415–421

Zhang R, Jia Z, Ma X, Ma H, Zhao Y (2020) Characterising the morphological characters and carbohydrate metabolism of oat culms and their association with lodging resistance. Plant Biol 22: 267–276

Zhang W jun, Li G hua, Yang Y ming, Li Q, Zhang J, Liu J you, Wang S, Tang S, Ding Y feng (2014) Effects of nitrogen application rate and ratio on lodging resistance of super rice with different genotypes. J Integr Agric 13: 63–72

Zheng X, Levine D, Shen J, Gogarten SM, Laurie C, Weir BS (2012) A high-performance computing toolset for relatedness and principal component analysis of SNP data. Bioinformatics 28: 3326–3328

Zuber U, Winzeler H, Messmer MM, Keller M, Keller B, Schmid JE, Stamp P (1999) Morphological traits associated with lodging resistance of spring wheat (Triticum aestivum L.). J Agron Crop Sci 182: 17–24

## Literature cited

Barrett JC, Fry B, Maller J, Daly MJ (2005) Haploview: analysis and visualization of LD and haplotype maps. Bioinformatics 21: 263–265

Berry PM, Sylvester-Bradley R, Berry S (2007) Ideotype design for lodging-resistant wheat. Euphytica 154: 165–179

Butler DG, Cullis BR, Gilmour AR, Gogel BJ, Thompson R (2017) ASReml-R reference manual version 4. Hemel Hempstead, HP1 1ES, UK

Gauch HG, Zobel RW (1990) Imputing missing yield trial data. Theor Appl Genet 79: 753–761

Gauch HG (1992) Statistical analysis of regional yield trials: AMMIanalysis of factorial designs. Elsevier, Amsterdam

Dahl A, Iotchkova V, Baud A et al (2016) A multiple-phenotype imputation method for genetic studies. Nat Genet 48:466–472

Dias CTD, Krzanowski W (2003) Model selection and cross validation in additive main effect and multiplicative interaction models. Crop Sci 43: 865–873

Gilmour AR, Thompson R, Cullis BR (1995) Average information REML: an efficient algorithm for variance parameter estimation in linear mixed models. Biometrics 5: 1440–1450

Luo MC, Tian CT, Li XJ, Lian JX (2007) Relationship between morpho-anatomical traits together with chemical components and lodging resistance of stem in rice (Oryza sativa L). Acta Bot Boreale-Occidentalia Sin 27: 2346–2353

Mackay TFC (1996) The nature of quantitative genetic variation revisited: Lessons from Drosophila bristles. BioEssays 18: 113–121

Paderewski J, Rodrigues PC (2014) The usefulness of EM-AMMI to study the influence of missing datapattern and application to Polish post-registration winter wheat data. Austral J Crop Sci 8: 640–645

Patterson HD, Thompson R (1971) Recovery of inter-block information when block sizes are unequal. Biometrika 58: 545

Pinthus MJ (1974) Lodging in wheat, barley, and oats: the phenomenon, its Causes, and Preventive Measures. Adv Agron 25: 209–263

Rodrigues PC, Malosetti M, Gauch HG, van Eeuwijk FA (2014) A weighted AMMI algorithm to study genotype-by-environment interaction and QTL-by-environment interaction. Crop Sci 54:1555–1570

Schindelin J, Arganda-Carreras I, Frise E, Kaynig V, Longair M, Pietzsch T, Preibisch S, Rueden C, Saalfeld S, Schmid B, et al (2012) Fiji: an open-source platform for biological-image analysis. Nat Methods 9: 676–682

Scutari M, Howell P, Balding D J, Mackay I (2014) Multiple quantitative trait analysis using bayesian networks. Genetics 198: 129–137

Wang B, Smith SM, Li J (2018) Genetic regulation of shoot architecture. Annu Rev Plant Biol 69: 437–468

